# A Spatiotemporal and Machine-Learning Platform Accelerates the Manufacturing of hPSC-derived Esophageal Mucosa

**DOI:** 10.1101/2023.10.24.563664

**Authors:** Ying Yang, Carmel Grace McCullough, Lucas Seninge, Lihao Guo, Woo-Joo Kwon, Yongchun Zhang, Nancy Yanzhe Li, Sadhana Gaddam, Cory Pan, Hanson Zhen, Jessica Torkelson, Ian A. Glass, the Birth Defects Research Laboratory, Greg Charville, Jianwen Que, Joshua Stuart, Hongxu Ding, Anthony Oro

**Author notes:** Correspondence should be addressed to H.D. or A.O.

## Abstract

Human pluripotent stem cell-derived tissue engineering offers great promise in designer cell-based personalized therapeutics. To harness such potential, a broader approach requires a deeper understanding of tissue-level interactions. We previously developed a manufacturing system for the ectoderm-derived skin epithelium for cell replacement therapy. However, it remains challenging to manufacture the endoderm-derived esophageal epithelium, despite both possessing similar stratified structure. Here we employ single cell and spatial technologies to generate a spatiotemporal multi-omics cell atlas for human esophageal development. We illuminate the cellular diversity, dynamics and signal communications for the developing esophageal epithelium and stroma. Using the machine-learning based Manatee, we prioritize the combinations of candidate human developmental signals for *in vitro* derivation of esophageal basal cells. Functional validation of the Manatee predictions leads to a clinically-compatible system for manufacturing human esophageal mucosa. Our approach creates a versatile platform to accelerate human tissue manufacturing for future cell replacement therapies to treat human genetic defects and wounds.

## INTRODUCTION

Directed differentiation from human pluripotent stem cells (hPSCs) hold great promise for regenerative and precision medicine^1^. Despite some progress, enormous hurdles in designing such differentiation strategies prevent rapid development and improvement of manufacturing protocols. State-of-the-art practices either recapitulate stepwise developmental processes revealed in model organisms, or rely on trial and error^2,3^. However, non-conserved developmental patterns between human and animal models impede the design of effective clinical strategies ^4, 5, 6^, while trial and error approaches are extremely labor-intensive and time-consuming. To address such limitations, a generalizable, unbiased approach is in demand to directly translate human tissue developmental signaling into actionable lineage-specific tissue manufacturing steps.

Recent advances in single cell and spatial technologies have revolutionized developmental and regenerative biology, bringing in unprecedented insights into tissue interaction signals for human development^7^. Emerging human developmental cell atlases, including gut, lung, heart and others, serve as an *in vivo* reference for the *in vitro* human cell engineering^8,9,10,11^. While these molecular maps provide a wealth of interactive signals, methodology to prioritize actionable and combinatorial inductive signaling for tissue manufacturing is lacking^12^.

We and others pioneered the derivation of genetically-corrected, hPSC-derived skin basal cells (BCs). BCs are tissue-specific stem cells capable of re-establishing the entire stratified epithelium, thus opening up possibilities for cell replacement therapy for genetic diseases like recessive-dystrophic epidermolysis bullosa (RDEB)^13,14,15^. RDEB patients also suffer from blistering of other stratified epithelia like esophagus and cornea^16,17^, but an equivalent therapy in esophagus, an endoderm rather than ectoderm derived tissue, has been challenging to translate. Previous studies have reported the derivation of esophageal progenitors by leveraging key signals revealed in mouse^18,19^. However, human and mouse esophageal epithelia show striking differences in tissue architecture and cellular activity^20,21^, suggesting discrepancies in their regulatory logic during development. Because of the extreme timing differences in gestation are reflected in pluripotent cell differentiation, mimicking mouse esophageal development to produce human esophageal mucosa has been unsuccessful. In practice, previous attempts resulted in inefficient human BC derivation that lacked current good manufacturing practice (cGMP) compatibility, underpinning the urgency to more deeply interrogate human esophageal development.

Underlying the enormity of the hurdle comes from our current limited knowledge of human esophageal development. Classic histology studies show that the human esophagus is specified in the dorsal anterior foregut and separated from the ventral trachea at around 4-6 weeks of human gestation^22^. At 8 weeks, the human esophagus becomes lined with stratified columnar epithelium that is protected by the terminal differentiation into multi-ciliated cells. Later at 20 weeks’ gestation, squamous stratified epithelium appears to replace the ciliated epithelium as the dominant terminal differentiation trajectory^23,24^. While the epithelial morphogenesis has been documented superficially, the underlying stromal composition and roles in morphogenesis remain unknown. Developmental studies in skin squamous stratified epithelium and our recent work have shown that regional mesodermal morphogenic signals wire the spatial-temporal specific stratification program in surface ectoderm progenitors, and these signals are required in our hPSC-derived skin production for optimal graftability^25,26^. Therefore, to accelerate esophageal mucosa manufacturing, a holistic molecular characterization of the coordinated development of epithelium and stroma is needed.

To overcome these hurdles, we have now built an integrated tissue informatics platform and present a spatiotemporal single-cell multi-omics atlas for human esophageal development. With such an atlas, we comprehensively catalog the cellular diversity, lineage transition, anatomic architecture and intercellular communication of the esophageal epithelium and stroma. We then use the novel deep learning algorithm Manatee to prioritize signaling pathway combinations for the *in vitro* specification of esophageal BCs (eBCs). We identify activating EGF, BMP, TGFb while inhibiting WNT as the optimal strategy for eBC derivation, and functionally validate the approach to establish a clinical standard hPSC-to-eBC differentiation system that will accelerate translational applications.

## RESULTS

### A spatiotemporal multi-omics cell atlas for human esophageal development

Aiming at manufacturing esophageal mucosa using human tissue developmental signaling, we comprehensively characterized the cellular dynamics during human esophageal development and extracted the intercellular signals associated with morphogenesis for *in silico* screening (Figure 1A). In collaboration with the NIH-sponsored UW Birth Defects Research Laboratory (BDRL), we ethically collected human embryonic and fetal esophagus samples from 35 individual donors for single cell and spatial profiling (Figures 1A-B, Table S1). Our dataset ranged from 45 (mid first trimester) to 132 (mid second trimester) post-conception days (noted as E45 to E132), spanning several key developmental milestones: onset of epithelial morphogenesis, fetal-specific ciliogenesis and squamous stratification (Figure 1B, Figure S2A). In addition to the epithelium, our dataset also catalogs the drastic growth, increased tissue complexity, and organization in the surrounding stroma (Figure 1B), thus providing the opportunity to explore tissue interactive signals during the coordinated development of epithelium and stroma.

**Figure 1.**
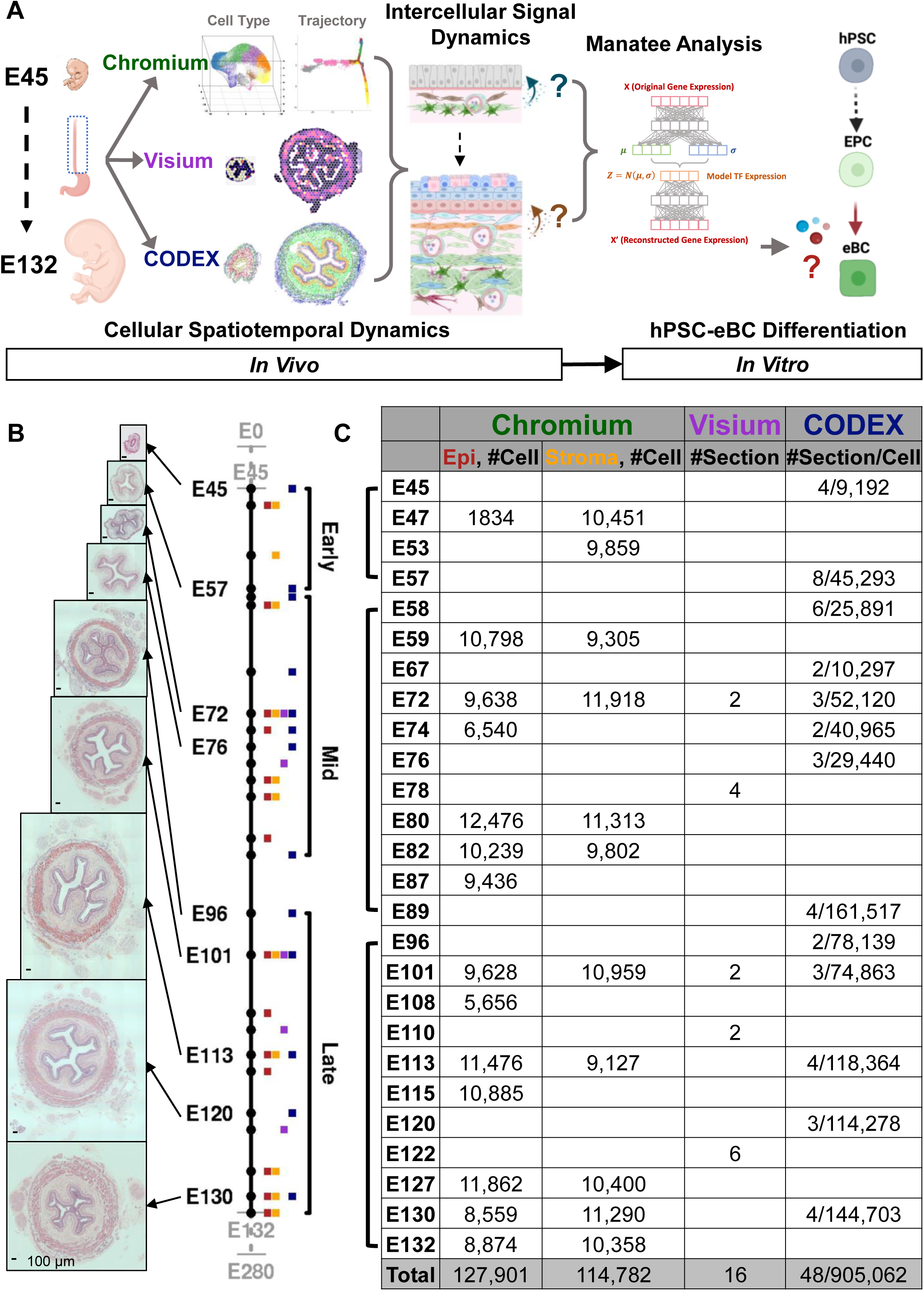
A multi-omics cell atlas of human esophageal development. (A) Workflow schematic: first integration of Chromium, Visium and CODEX to build the multi-omics cell atlas to depict human esophageal development, followed by investigation into intercellular signal dynamics to nominate BC specification signals. Then the deep-learning algorithm Manatee predicts the optimal signal combination to drive BC specification, which is further used to build the hPSC-eBC manufacturing platform. (B) H&E staining of developing human esophageal sections from E45 to E130 with the timeline and summary of analyzed samples colored by assays. Scale bars: 100 μm. (C) Summary showing the total number of cells collected in each dataset.

By densely sampling human esophageal development, we produced high quality single cell transcriptome data for both epithelium and stroma using distinct cell dissociation methods (Figure 1C, Figures S1A-B). After iterative clustering, removal of low-quality and contaminant clusters, and removal of the erythrocyte cluster, we present 97,212 epithelial cells from 14 timepoints and 91,891 stromal cells from 11 timepoints (Figures S1B-D). Using canonical markers, we grossly annotated 6 major cellular compartments: epithelium (EPI), mesenchyme (MES), enteric nervous system (ENS), endothelium (ENDO), skeletal muscle (SKM), immune cells (IM). Fine-grained clustering of each compartment further discovered 10 cell types/states in EPI (Figure 2A, Figures S1B, S2B-D) and 66 in STROMA (Figure 3A, Figures S1B, S2E-G, S3).

**Figure 2.**
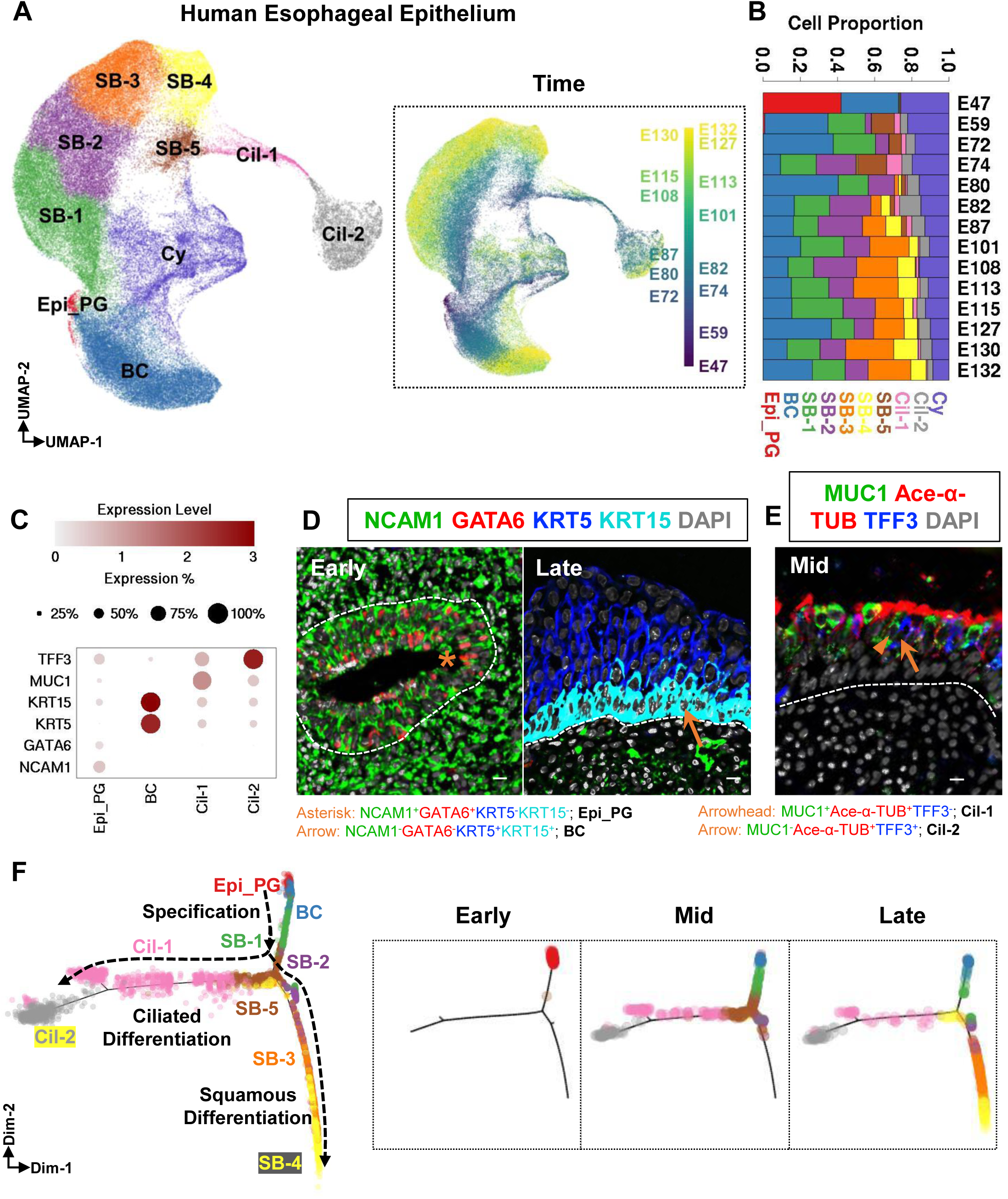
The cellular heterogeneity and lineage trajectory of human esophageal epithelium during development. (A) Epithelial single cells were projected on the UMAP space, and color-coded based on cell type. Inset showing UMAP visualization by developmental timepoint. (B) Proportions of the identified cell types at each timepoint. (C) Dot plot showing the expression levels for selected markers of Epi_PG, BC, Cil-1 and Cil-2 shown in CODEX images. (D-E) CODEX images confirming the presence of Epi_PG, BC (D), Cil-1, Cil-2 (E) using selected markers. Early: Representative images from E45 (Early), E120 (Late) and E72 (Mid) samples. Scale bars: 10 μm. (F) Monocle2 trajectory of the whole epithelial compartment, color-coded based on cell type. Insets: projection of epithelial cells collected at representative stages on the Monocle2 trajectory. Early: E47; Mid: E72; Late: E130.

**Figure 3.**
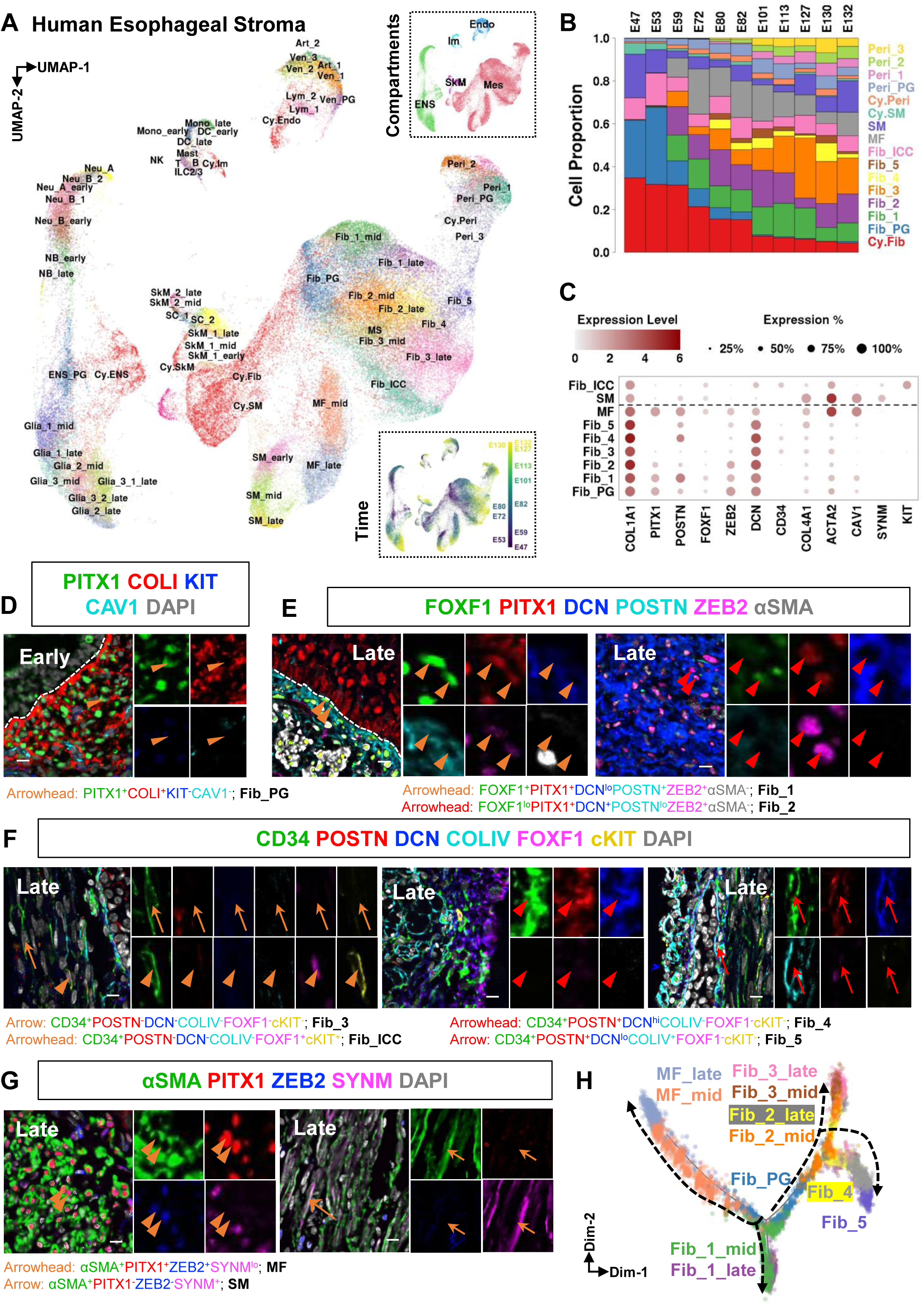
Esophageal stromal diversity and mesenchymal lineages. (A) Stromal single cells were projected on the UMAP space, and color-coded based on cell types/states. Inset showing UMAP visualization by cellular compartment (top right) and developmental timepoint (bottom right). (B) Proportions of the identified cell types at each timepoint. (C) Dot plot showing the expression levels for selected markers of different fibroblasts, MF and SM shown in CODEX images. (D-G) CODEX images confirming the presence of Fib_PG (D), Fib_1/2 (E), Fib_3/4/5, Fib_ICC (F), MF/SM (G) using selected markers. Early: Representative images from E45 (Early) and E120 (Late) samples. Scale bars: 10 μm. (H) Monocle2 trajectory of the Fib-MF lineages, color-coded based on cell type.

Classic tissue recombination experiments demonstrate that the signal interactions between epithelium and local mesenchyme are essential for tissue morphogenesis and cell fate determination^27^. We and others have demonstrated the significance of integrating supporting cellular signals into human tissue engineering^25,26, 28^, reinforcing the importance of detailing the spatial architecture of developing human esophagus. To systematically map the cell types revealed in single cell atlas, we orthogonally performed unbiased Visium spatial transcriptomics (16 sections from 5 timepoints, 5 donors) and high-resolution CODEX (co-detection by indexing) proteomic profiling (48 sections/905,062 cells from 13 timepoints, 15 donors) (Figure 1C, Table S1). We integrated single cell transcriptomics with the spatial profiles to delineate the dynamic cellular architecture for human esophageal development. The full dataset can be explored online through the EsoDev data portal (http://visualify.pharmacy.arizona.edu/EsoDev/).

### Developmental origin of BCs identified at the early stage

We first cataloged human esophageal epithelial morphogenesis to build the molecular roadmap for esophageal mucosa manufacturing, focusing on the major tissue morphogenic transitions. Our scRNA-seq dataset of human esophageal epithelial cells showed a relatively constant proportion of KRT5^+^/KRT15^+^ BCs across all the samples, suggesting the establishment of the self-renewing BC stem cell pool in early development (Figures 2A-B, Figures S2B-D). To identify BC precursors, we focused on the small cluster of cells (Epi_PG) negative for BC or any other known differentiated cell markers. These cells were only detected in the earliest E47 scRNA-seq sample (Figures 2A-C, Figures S2B-D). Their high expression of SHH and GATA6 identified the uncommitted foregut progenitor identity^29,30,31^. Their presence was validated by immunofluorescence (IF) as NCAM1^+^/GATA6^+^ in early-stage esophageal sections before the appearance of KRT5^+^/KRT15^+^ BCs (E45 shown in Figure 2D, E57 & E58 data not shown). By H&E, we found that these cells presented a primitive-appearing morphology in the multilayered early esophagus (Figure 1B, Figure S2A, E45 and E57). Furthermore, the Monocle pseudotime analysis predicted that Epi_PG first give rise to BCs, while BCs later generate other differentiated populations (Figure 2F). Therefore, for the first time we reveal the molecular features and lineage commitment of the stratified columnar epithelium documented in classic histology literatures.

### Two differentiation waves of esophageal epithelial morphogenesis

After the specification of BCs, we observed two distinct differentiation programs that are sequentially activated: fetal-specific ciliated differentiation and canonical squamous stratified differentiation (Figures 2F, 4C). The ciliogenesis trajectory originates from BCs, transitioned through first KRT15^-^/KRT5^+^/KRT13^+^/KRT4^-^ suprabasal cell (SB-1) and then SCGB1A1^+^/UPK1B^+^ SB-5 to FOXJ1^+^/Ace-α-TUB^+^ ciliated cells (immature MUC1^+^/TFF3^-^ Cil-1 and mature MUC1^-/^TFF3^+^ Cil-2, Figures 2E-F, Figures S2B-D). By contrast, the squamous stratification trajectory follows BCs, SB-1, and then KRT15^-^/KRT5^lo^/KRT13^+^/KRT4^lo^ SB-2, KRT15^-^/KRT5^-^/KRT13^+^/KRT4^lo^ SB-3 and KRT15^-^/KRT5^-^/KRT13^+^/KRT4^+^ SB-4 (Figure 2F, Figures S2B-D). Interestingly, the ciliogenesis was predominantly observed at mid stage, marked by the peak of SCGB1A1^+^/MSLN^+^/KRT4^+^ SB-5 before E80 (Figure 2B). As SB-5 is the only cluster that serves to replenish ciliated populations, the exhaustion of SB-5 by E101 underlies the later gradual lineage extinction. The alternative canonical squamous stratification starts from E80, marked by the appearance of SB-3 and SB-4. The rapid increase in the proportions of SB-3/4 after E101 represents the replacement of ciliated differentiation by the ultimately dominant squamous stratification (Figures 2B, 4C). Thus, we depict two conversion events in epithelial morphogenesis: the primitive multilayered epithelium at the early stage first differentiating to ciliated columnar epithelium and marked by specification of BCs and ciliated differentiation; later during the mid stage, the squamous stratified epithelium program replaces the transitional ciliated columnar epithelium and further matures the mucosa at later stages.

**Figure 4.**
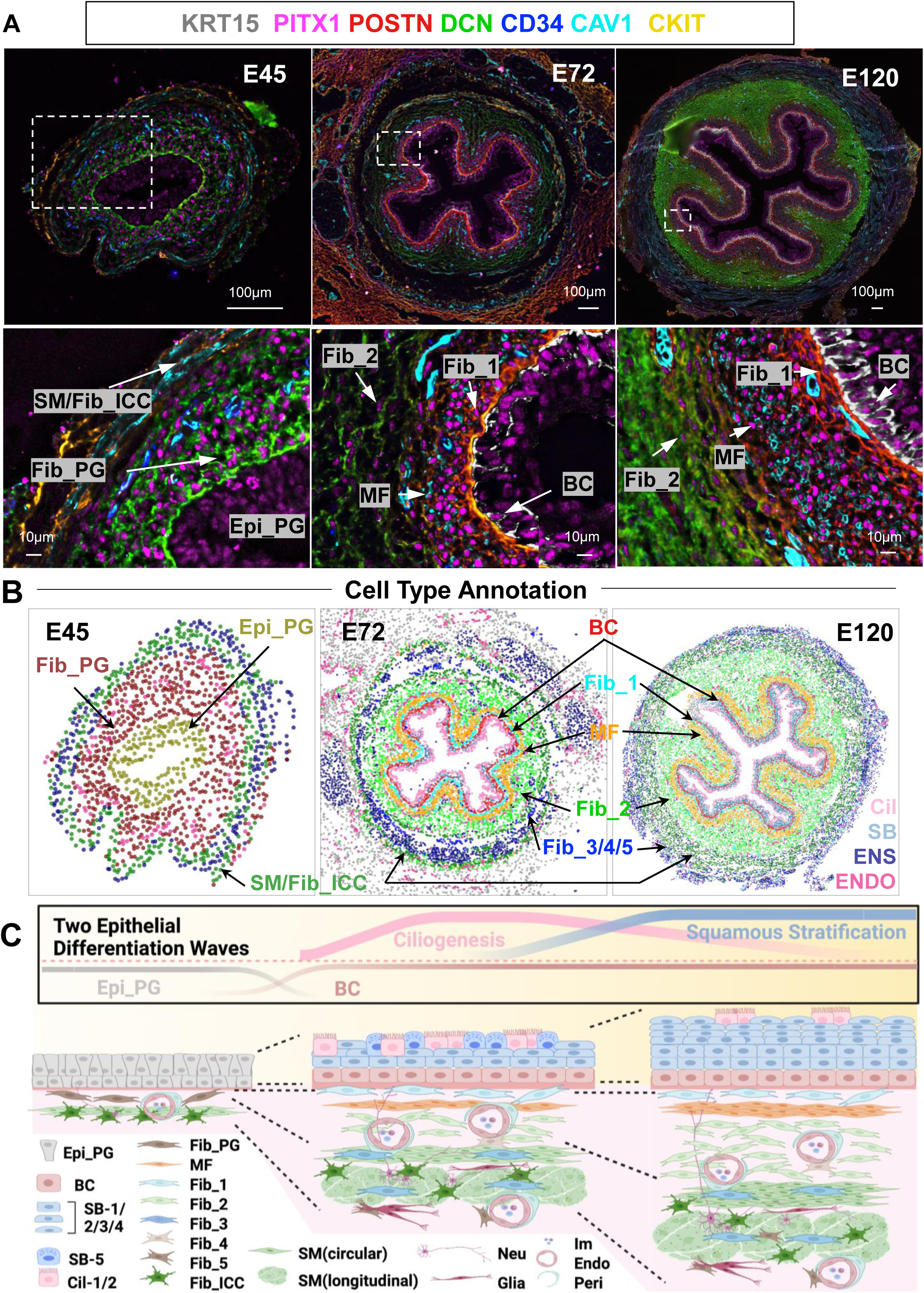
Spatiotemporal cellular dynamics of human esophageal development. (A) Representative CODEX images with selected markers showing the epithelium and the local mesenchyme at E45 (Early), E72 (Mid) and E120 (Late). Scale bars: 100 μm. Bottom panel showing enlarged boxed regions with representative cells annotated. Scale bar: 10 μm. (B) Cell type annotation of the representative CODEX images based on hierarchical clustering analysis of the fluorescent intensities of 42 markers. (C) Diagram of the cellular composition and tissue architecture at early, mid, late stages of the developing human esophagus.

### Myofibroblasts and fibroblasts share a common developmental origin

To interrogate the accompanying mesenchymal dynamics with these two epithelial differentiation waves, we subclustered the MES compartment and revealed 23 cell types/states. Like the overlying epithelium we detect a fibroblast progenitor cluster (Fib_PG, COLI^+^/PITX1^+^/ZEB2^+^/SNAI2^+^) that was prominent at the early stage (E47 and E53), but lacking at later stages (after E101) (Figures 3B-D, Figures S2E-G). As Fib_PG abundance decreases from E59, we observe the emergence of three major COL1A1^+^ fibroblast subtypes Fib_1/2/3 and 1 myofibroblast population MF. The Fib_1 cells (FOXF1^+^/PITX1^+^/DCN^lo/-^/POSTN^+^/ZEB2^+^) express high levels of POSTN and TNC, suggesting a potential regulator role in epithelial proliferation and differentiation through paracrine signaling (Figures 3C, 3E, Figures S2G, S4A-B)^32,33,34,35^. The Fib_2 cells (FOXF1^lo/-^/PITX1^+^/DCN^+^/POSTN^lo/-^/ZEB2^+^) express high levels of collagen (COL1A1, COL3A1, COL14A1), elastic fiber (FBLN1, FBLN5, FBN1, FBN2) and proteoglycan (DCN, LUM) genes, all of which could provide structural support for the esophagus (Figures 3C, 3E, Figures S2G, S4A-B)^36,37,38^. Fib_3 cells (CD34^+^/POSTN^-^/DCN^lo/-^/COLIV^-^/FOXF1^lo/-^) specifically express the calcium-activated potassium channel gene KCNN3, the sodium channel gene SCN7A and the CD34 marker, suggesting their potential roles in neurotransmission and regulation of muscular motility (Figures 3C, 3F, Figures S2G, S4A-B) ^39, 40^ . MF cells (αSMA^+^/CAV1^+^/DES^+^/PDGFRA^+^) share common markers with Fib_1, including PITX1, ZEB2, DPT, POSTN and TNC (Figures 3G-H, Figures S2E-G, S4A-B), suggesting potential roles in regulating epithelial regeneration. Two additional fibroblast subtypes, Fib_4/5, were found only after E80, suggesting a developmental delay in their specification, and possibly a later role in tissue functions. Indeed, Fib_4 cells (CD34^+^/POSTN^+^/DCN^+^/COLIV^-^/FOXF1^-^) express genes related to angiogenesis and vascular contractility (ANGPTL1 and PRRX1), underlying a putative perivascular fibroblast identity (Figures 3C, 3F, Figures S2G, S4A-B) ^41, 42^ . Fib_5 cells (CD34^+^/POSTN^+^/DCN^lo/-^/COLIV^+^/FOXF1^-^) exclusively express the PI16 gene and therefore considered to be adventitial fibroblasts ^43, 44^ . Interestingly, Fib_5 cells also express multiple glia-related genes (DCX, PLP1, NGFR and ENTPD2), suggesting a close relationship with the ENS compartment (Figures 3C, 3F, Figures S2G, S4A-B). The pseudotime trajectory analysis predicts that Fib_PG first differentiates into three branches, Fib_1, Fib_2, and MF; while Fib_2 further follows two branches, one into Fib_3 and another into Fib_4/5 (Figure 3H).

### Interstitial cells of Cajal and smooth muscle lineages are specified early in development

The highly specialized mesenchymal cells, interstitial cells of Cajal (Fib_ICC, CD34^+^/cKIT^+^/ETV1^+^/ANO1^+^), and the smooth muscle cells (SM, αSMA^+^/SYNM^+^/CAV1^+^/DES^+^) are essential executors for the rhythmic peristalsis of esophagus^45,46,47^. These two cell types were constantly found in all the samples across the developmental stages (Figures 3B-C, 3F-G, Figures S2E-G). Consistent with the signaling coordinator role between enteric nerves and muscles, Fib_ICC was found intermingled with SM (Figures 4B-C)^48^. Both cell types followed their isolated maturation trajectories after specification before E47, thus revealing an early establishment of the contractile muscular components. We did not observe common progenitors for Fib_ICC and SM, or any potential transdifferentiation between these two lineages^49,50,51,52^. Even though subpopulations of Fib_ICC and SM have been described based on their anatomical locations, they were both found as relatively homogenous populations that mature over time^53^.

### Pericyte diversity is established with a developmental lag

Another key component in the MES compartment is the pericyte (Peri, PDGFRB^+^/RGS5^+^/NOTCH3^+^) (Figure 3A, Figures S2E-G). αSMA^+^/NG2^+^ pericytes were visualized surrounding vasculature (Figures S3L-M). In contrast to the isolated lineages of Fib_ICC and SM, Peri exhibits much more diversity and delayed development. The progenitor population, Peri_PG, were observed as early as E47. This population maintained a relatively stable proportion throughout the subsequent stages, thus potentially serving as a self-maintaining progenitor pool. From E80, we detected three other pericyte populations, differentiating (Peri_1, THBS1^+^), contractile (Peri_2, ACTA2^+^/MYL9^+^/MYH11^+^) and angiogenic (Peri_3, PRRX1^+^/PROCR^+^) pericytes (Figures 3A-B, Figures S2E-G)^54,55^. Notably, Peri_3 and Fib_4 were partially overlapped on UMAP, reflecting their common features in angiogenic genes (Figures S2E-G). However, we could not find evidence supporting lineage transformation between Peri and MF, which has been suggested in some fibrosis studies^56^.

### Neurogenesis precedes gliogenesis to shape the enteric nervous system

ENS is the second largest compartment in esophageal stroma, including 16 distinct clusters (Figure S1B). Enteric neural crest cells first enter esophagus and migrate caudally to colonize the entire gut tube^57^. Consistent with previous literature, we observed abundant progenitor population (ENS_PG, SOX10^+^/SOX2^+^/S100B^+^/TUBB3^+^/PGP9.5^+^) in E47 and E53 samples (Figure 3A, Figures S3A-E). ENS_PG quickly track the TUBB3^+^/UCHL1^+^ neuronal differentiation path via neuroblasts (NB, ASCL1^+^/DLL3+). NB cells maintain a progenitor pool while continuously giving rise to the ETV1^+^ Neuron_A (Neu_A) and BNC2^+^ Neuron_B (Neu_B) populations (Figures S3D, S3G)^58^. The presence of Neu_A and B at E47 suggested an early neurogenesis (Figure S3B). Meanwhile, ENS_PG cells give rise to the S100B^+^ esophageal glial lineage (Figure S3F). We identified 3 glial cell types, including BCAN^+^/APOE^+^/ENTPD2^+^ Glia_1, RXRG^+^/GFRA2^+^ Glia_2 and MPZ^+^/MAL^+^/DHH^+^ Glia_3. The glial cells are enriched in later stages compared to the neuronal lineage. Glia_1 was first observed at E59, while Glia_2 and 3 were detected only after E80 (Figure S3B). Interestingly, the ENS differentiation pattern described here in human developing esophagus matches well with the one revealed in human developing intestines^58^, suggesting that the gastrointestinal ENS system undergoes a coordinated differentiation program despite differences in progenitor types, colonization time and migratory paths^59^.

### Esophageal endothelial cells, skeletal muscles and immune cells establish cellular heterogeneity in early development

We also discovered esophageal ENDO, SKM and IM populations, which together comprise ∼10% of our STROMA cell collection (Figure 3A, Figure S1B). We highlighted an unexpected early establishment of cellular heterogeneity in these compartments. For ENDO, capillary (Art_1, CXCR4^+^/PGF^+^/LXN^+^) and large (Art_2, HEY1^+^/GJA5^+^) arterial cells, as well as lymphatic vessel cells (Lym_1) were observed as early as E47. Lymphatic valve cells (Lym_2, SCG3^+^/FOXC2^+^/GATA2^+^) emerge at E80 (Figures S3H-K)^9,60,61,62^. This suggest that the establishment of arterial and lymphatic lineages occur earlier than E47. Notably, while the venous progenitors (Ven_PG) were also observed at E47, their fraction peaks at E53, and becomes marginal at E80. Simultaneously, the potential intermediate venous cells (Ven_1) are being specified. Subsequently, large (Ven_2, ACKR1^+^/ADGRG6^+^) and capillary (Ven_3, CD83^+^/RGCC^+^/CA4^+^) venous cells emerge, which were first observed at E72^63^. These observations delineate the time window of venous lineage commitment during human esophageal development (Figures S3H-M).

For SKM, we identified PAX7^+^/MYF5^+^ SkM_1 and MYOG^+^/MYF6^+^ SkM_2 skeletal muscle cells,as well as SOX8^+^/PAX7^lo^/MYF5^lo^ SC_1 and SOX8^lo^/PAX7^+^/MYF5^+^ SC_2 putative satellite cells^64,65,66^. These four populations were observed at all developing stages, suggesting early specification during esophageal myogenesis (data not shown). Meanwhile, such an entire skeletal muscle lineage expresses pharyngeal mesoderm transcription factors (ISL1, TBX1, MSC and SIX1) and lacks PAX3, which is consistent with the non-somitic, cranial mesodermal origin previously revealed in mouse studies^64,67^. Finally, we found the presence of 6 IM populations as early as E47, a surprising finding given the lack of environmental antigen exposure. CD45^+^ IM cells were found scattered among epithelium and stroma (Figure S3N). Identified populations included monocytes (Mono, CD163^+^), dendritic cells (DC, LYZ^+^ or HLA-DRA^+^), mast cells (Mast, KIT^+^/TPSB2^+^) from the myeloid lineage, and B cells (B, IGHM^+^/CD79A^+^/CD79B^+^), natural killer cells (NK, GZMA^+^/KLRD1^+^) and type 2 and 3 innate lymphoid cells (ILC2/3, GATA3^+^/RORA^+^/RORC^+^/KIT^+^) from the lymphoid lineage ^68, 69, 70^ . T cells (T, CCR7^+^/CD3D^+^/CD3G^+^) emerge at E80, revealing the later migration from the thymus to the developing esophagus^71^, and emphasizing the presence of the innate rather than adaptive immune system during esophageal development (data not shown).

### Spatiotemporal cellular dynamics establish esophageal tissue architecture

With the cell types identified, we orthogonally performed CODEX and Visium to dissect the spatiotemporal orchestration of developing esophageal epithelium with its surrounding stroma. Based on the single cell gene expression pattern, we designed a CODEX panel targeting 42 markers. Such markers, on their own and in combinations, uniquely identify the majority of discovered cell populations. With such a panel, we performed CODEX profiling, further determined cellular identities by combining hierarchical clustering and marker-guided annotation (Figure S4, see METHODS). Without losing generalizability, we selected one representative CODEX section per developmental time window for a detailed spatiotemporal survey. As shown in Figures 4A-B, the early window (E45) of columnar epithelium features a less differentiated esophagus rudiment: Epi_PG is the only cell type in the epithelium, and Fib_PG dominates the surrounding inner stromal layer. The mid window (E72), marks coordinated differentiation processes in epithelial and mesenchymal compartments. The appearance of BC coincides with mesenchymal stratification formed by Fib_1, MF, Fib_2, Fib_3/4/5 and SM/Fib_ICC layers. Fib_1 and MF closely surround the basement membrane. While Fib_3/4/5 intermingle with Fib_ICC/SM in the muscularis propria, Fib_2 cells are located between MF and the muscularis propria. The late window (E120) presents a continuous increase in the size of the developing esophagus. Additionally, we noticed that ENS neurons and glia cells aggregate in the adventitial plexus and myenteric plexus. By contrast, ENDO cells and IM cells scatter with no noticeable spatial aggregation.

We further unbiasedly confirmed the spatial distribution of the major cell types at mid and late stages by integrating scRNA-seq and Visium profiles using cell2location, (see METHODS)^72^. Additionally, Neu and Glia cells exhibited distinct spatial enrichment. Neu cells were more scattered with muscularis propria, while Glia tended to aggregate in adventitial plexus (Figure S5). In summary, our multi-omics atlas depicts the detailed spatiotemporal cellular dynamics of human esophageal development. At early stage, the esophageal rudiment is lined with progenitor-only epithelium which is surrounded by fibroblast progenitors and primitive muscularis propria. Along with BC specification, these fibroblast progenitors differentiate into diverse lineages and establish the highly organized stromal architecture. The stromal cells further mature and expand from mid to late stage when the squamous differentiation wave replaces the ciliogenesis wave in the epithelial compartment (Figure 4C).

### Spatiotemporal candidate signal nomination inspired by the multi-omics atlas

Following delineating cellular dynamics of human esophagus development, we focused on local epithelium-mesenchyme signaling interactions driving eBC development, which could be leveraged to manufacture eBCs from hPSCs (Figure 5A). We noted that the Fib_PG, SM and Fib_ICC are the local mesenchyme for Epi_PG in the early stage, and Fib_1, Fib_2, MF for eBCs in mid and late stages based on their spatial distribution in Visium and CODEX analyses (Figure 4, and Figure S5). Importantly, Fib_1 maintains a constant 10 µm cellular layer adjacent to eBCs throughout the developmental process immediately surrounded by the MF layer that doubles its thickness only at the late stage. By contrast, layers Fib_2 and SM/Fib_ICC proliferate and thicken dramatically during development, which increases the overall distance from the eBCs (Fib_2 is located 69.70 ± 4.40 µm from the basement membrane at E58, 103.57 ± 9.51 µm at E72, 223.37 ± 21.26 µm at E120. SM/Fib_ICC is located 104.50 ± 6.74 µm from the basement membrane at E58, 219.65 ± 14.68 µm at E72, 497.53 ± 37.66 µm at E120) (Figure 5B). Considering the effective range of common morphogen gradients^73,74,75^, we conclude that signals emanated from SM/Fib_ICC are less likely to strongly affect eBCs after the early stage. Fib_1, Fib_2 and MF, on the other hand, emerge as key signal senders at mid and late stages.

**Figure 5.**
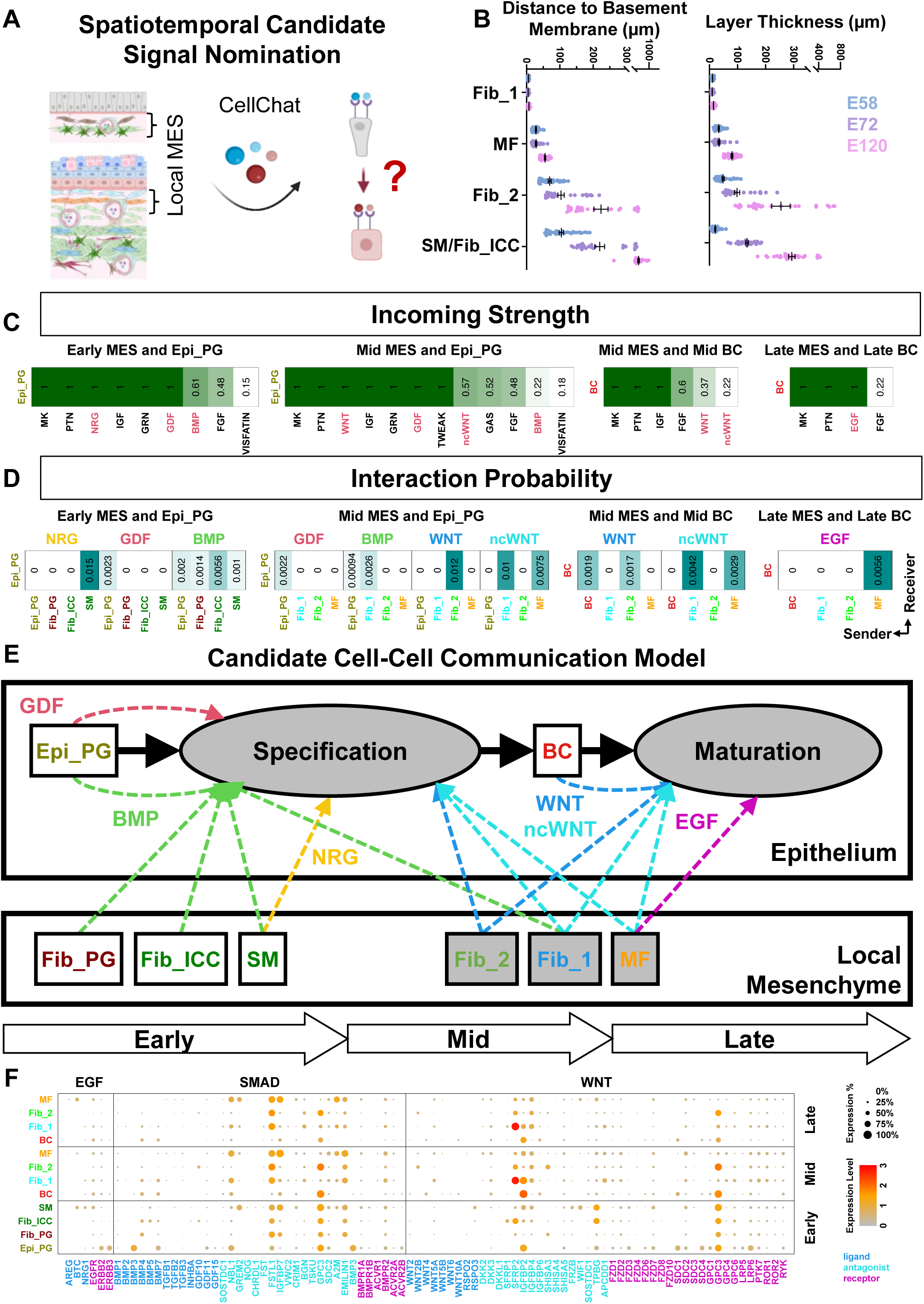
Spatiotemporal candidate signal nomination. (A) Schematic: determining local mesenchymal cellular components at different developing stages first, followed by identifying potential inductive signals from the local mesenchyme that could be received by Epi_PG/BC using CellChat. (B) Quantification of spatial distance to basement membrane (left panel) and layer thickness (right panel) of key mesenchymal cell types. Boxplots represent mean ± SEM. Each dot represents one random measurement. For each stage, n >= 30. (C-D) Heatmaps showing the CellChat predicted incoming signal strength received by Epi_PG/BC (C) and sender-receiver interaction probability (D). (E) Diagram summarizing the candidate intercellular communication model of esophageal BC development. (F) Signaling gene expression patterns of key signaling pathways summarized in a dot-plot. For each pathway, expression patterns of ligand, antagonist and receptor genes among early, mid and late stage cell populations were visualized. The expression level and percentage were coded by dot color and size, respectively.

Next, we used the CellChat algorithm to predict stage-dependent signaling interactions between the local mesenchyme and Epi_PG/BC in an unbiased manner^76^. As the Epi_PG-to-BC specification is a continuous developmental process, we reasoned that both early and mid local mesenchymal components would be sources for inductive signals. Therefore, we first characterized signals that are received by Epi_PG and BC in early and mid stages, and we identified NRG, GDF, BMP, WNT and ncWNT ranking as top candidates (Figure 5C). Meanwhile, we associated BC maturation with the mid-late local mesenchyme, and found EGF, WNT and ncWNT as inductive signals.

We further interrogated the cellular sources of the above signals. For BC specification, at the early stage, SM is the only sender for NRG, GDF is autocrine by Fib_PG, while Fib_ICC serves as the major provider for BMP. At the mid stage, SM no longer serves as the local mesenchyme which causes the loss of NRG. Fib_1 replaces Fib_ICC as the major source for BMP. Fib_1, together with other emerging populations including Fib_2 and MF, introduce WNT and ncWNT which were not observed before. Fib_1, Fib_2 and MF continue providing WNT and ncWNT for BC maturation at the mid stage. The BC serves as additional source for WNT in an autocrine manner. At the late stage, the only signal is EGF, which is secreted solely by MF (Figure 5D).

The above results suggest a dynamic signal communication pattern alongside the BC specification-maturation trajectory (Figure 5E). We further confirmed cellular signal sender/receiver identities with single cell expression patterns of known ligands, agonists and antagonists (Figure 5F). Taken together, we concluded that EGF superfamily (EGF and NRG), WNT pathway (WNT and ncWNT), TGFb superfamily member GDF and BMP pathway might serve as key inductive signals to drive BC development.

### Manatee facilitates design of hPSC-eBC differentiation strategies

Manufacturing eBCs *in vitro* for clinical applications remains challenging as the latest protocols produce progenitors rather than lineage-specified, functional eBCs, and are also not chemically-defined^18,19^. While our atlas reveals the dynamic local mesenchymal signals which could be received by Epi_PG/BC and potentially drive BC development, the presence of both ligands, agonists and antagonists obscures whether those signaling pathways are activated or inhibited, and which signaling pathways cooperate with each other. The machine-readable nature of the integrated atlas data, in combination with the rapid development of more complex computational models allowed us to take an unbiased approach in prioritizing signaling combinations. We thus developed the Manatee deep learning framework to screen *in silico* for the optimal combination of EGF, WNT, TGFb and BMP signals for eBC specification.

Manatee predicts gene expression alterations caused by transcription factor (TF) perturbations. As shown in Figure 6A, our design was adapted from the variational autoencoder (VAE) model ^77^ . Specifically, we constrained the VAE latent space to represent TF expression, by adding an additional TF reconstruction loss during training. As for the prediction phase, we adjusted latent variable values associated with candidate TFs to be perturbed. The adjusted latent space will further be flowed through the decoder neural network. The outputs will be considered as the altered expression profiles, which is ensured by the generative capacity of VAEs (see METHODS).

**Figure 6.**
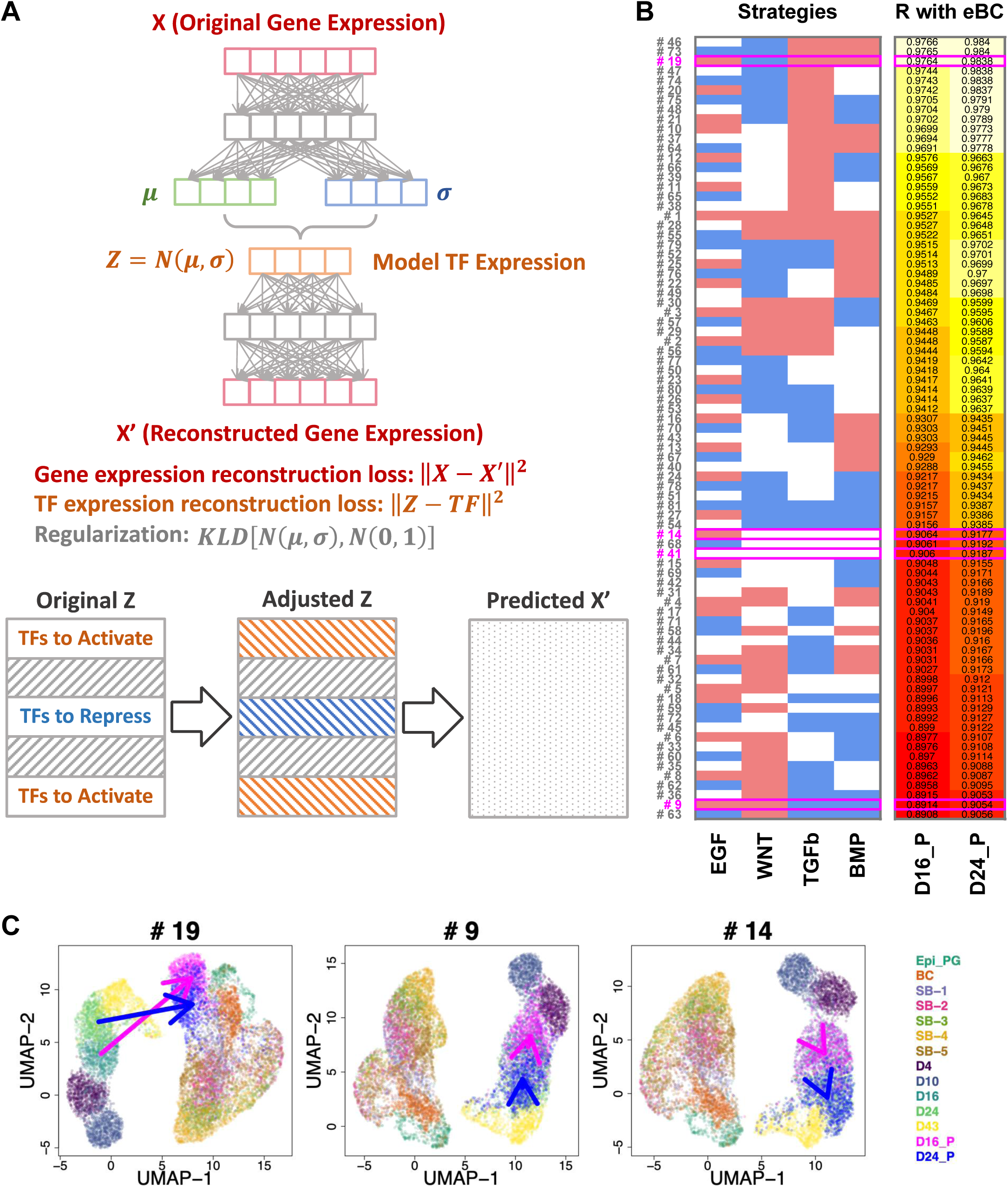
Manatee screens for the optimal signal combinations to promote eBC derivation. (A) The Manatee workflow. The Manatee model was adapted from VAE by constraining its latent space to represent TF expression. To predict perturbation-induced expression profiles, latent variables associated with candidate TFs will be adjusted. Such an adjusted latent space will then be decoded as final predictions. (B) Screening all 81 possible perturbation combinations of EGF, WNT, TGFb and BMP signaling pathways for eBC derivation (red, up-regulate; blue, down-regulate; white, intact). The perturbation effect was quantified by computing the Pearson correlation (R) between the average eBC and the perturbation expression profiles. Specifically, 4 combinations were highlighted, including the candidate optimal combination #19, negative control #9, base control #14 and #41 which has not been perturbed. (C) Original single cell expression profiles and predictions yielded by the above-highlighted strategies were visualized on the two-dimensional UMAP space. D4, 10, 16, 24 and 43, single cells harvested at day 4, 10, 16, 24 and 43 from the hPSC culture, respectively. D16_P and D24_P, predicted single cell profiles with D16 and D24 as starting points, respectively. Epi_PG, BC, SB-1 to 5, epithelium single cell populations as described in Figure 2. Arrows indicate perturbation effects.

We used Manatee to screen all 81 combinations of EGF, WNT, TGFb and BMP (Figure 6B), and evaluated the similarity between virtually perturbed cellular statuses and *in vivo* human developing BCs. We first derived esophageal progenitors from hPSCs using a new chemically-defined, xeno-free protocol^18,19,78^ (Figure 7A). Our bulk RNA-seq analysis of hPSC differentiation confirmed endoderm/foregut commitment from D4 to D10, whereas esophageal progenitors were specified by D16. These progenitors matured into eBCs after another month (Figures S7A-D). We then trained Manatee with D4, D10, D16, D24, D43 *in vitro* derived single cells (Figures S6A-C), together with *in vivo* epithelial cells as reported in Figure 2A, to capture the complete set of regulatory logics related to the eBC development. We further performed predictions with D16 and D24 cells as starting points (see METHODS). We quantified the eBC derivation effectiveness with the Pearson correlation between the average *in vivo* human developing BC and the perturbation expression profiles (Figure 6B). Considering that EGF is empirically beneficial for epithelial cultures, we focused on strategies with EGF activation. Based on the readout in effectiveness, strategy #19 was prioritized (it also ranked the 3rd best overall). Its reverse strategy #9, compared to the no-perturbation group #41, shows a notable decrease in effectiveness. Strategy #14 which has only EGF activation shows no obvious gain in the effectiveness compared to #41. We included strategies #19, #9 and #14 for experimental validations, as experimental, negative control and base control groups respectively. We also confirmed the necessity for pathway perturbations in deriving eBCs. We prolonged the culture to day 43 (D43) without any additional perturbation recipe and performed single cell profiling. The effectiveness of D43 culture score 0.9351, which is lower than our best combination #19 (0.9764 for D16_P, 0.9838 for D24_P). Meanwhile, UMAP dimension reduction suggests that #19 drives D16/24 cells to the eBC lineage, whereas D43 cells still track the *in vitro* cell culture trajectory (Figure 6C and Figure S6D).

**Figure 7.**
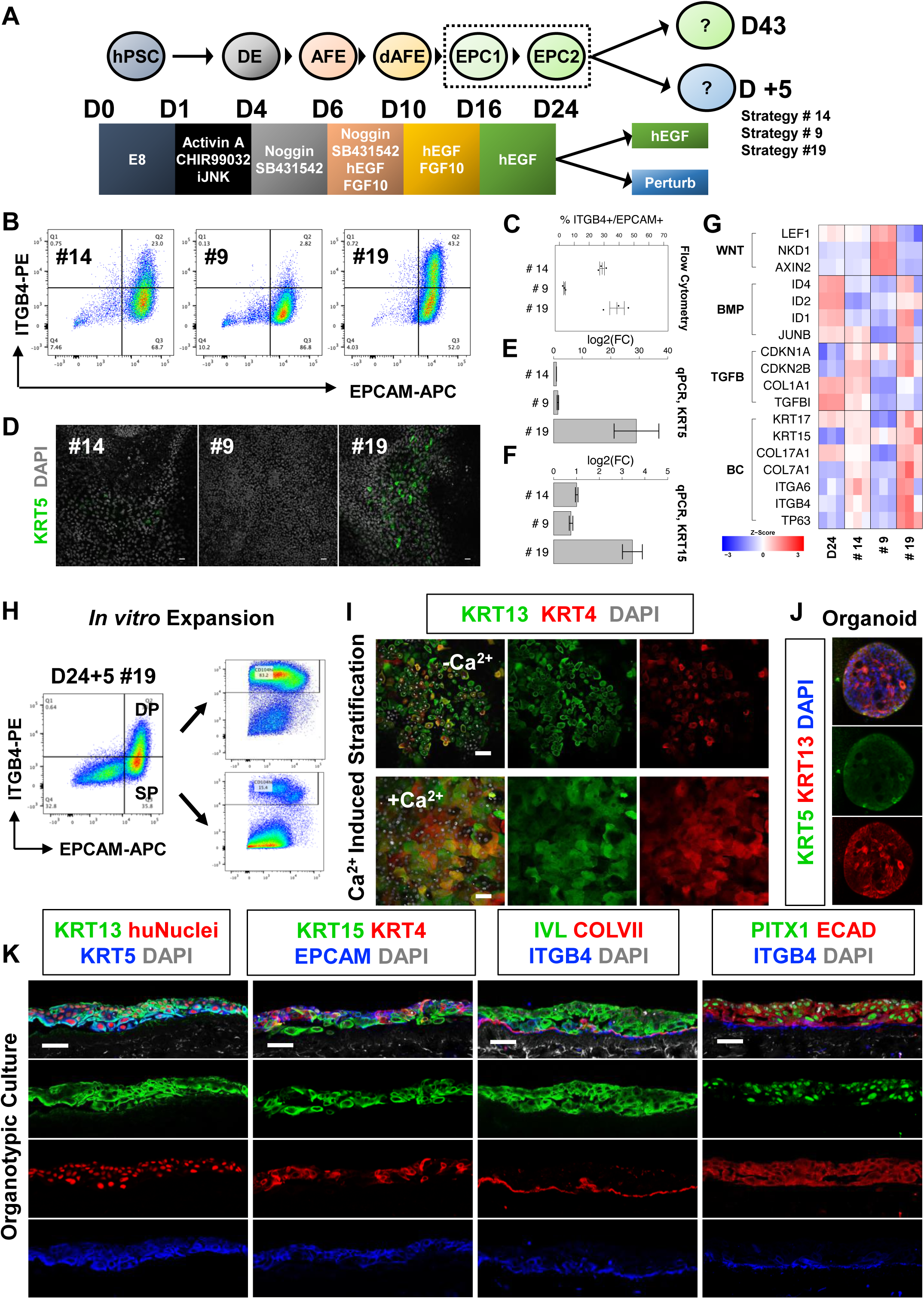
Establishment of eBC manufacturing platform using the Manatee-predicted strategies. (A) Schematic illustration of the hPSC-to-eBC differentiation protocol. DE, definitive endoderm; (d)AFE, (dorsal) anterior foregut endoderm; EPC, esophageal progenitor. EPCs were further subjected to a prolonged culture till D43 or a 5-day perturbation. (B) Representative flow cytometry plots of cells after the 5-day perturbation. Percentage of candidate eBCs (EPCAM^+^/ITGB4^+^) among all cells were quantified in strategy #19 (43.2%), negative control #9 (2.82%) and base control #14 (23%). (C) Statistical quantification of the flow cytometry analysis. Candidate eBC percentage was calculated against epithelial cells (EPCAM^+^). Boxplots represent mean ± SEM; n = 3. (D) IF staining of cells after the 5-day perturbation with general BC marker KRT5. (E, F) qPCR analysis of general BC markers KRT5 and KRT15. Marker expression was quantified by log2 fold change against base control #14. Boxplots represent mean ± SEM; n = 3. (G) Bulk RNA-seq heatmap of D24 cells before treatment and D29 cells after 5-day perturbation. Expression of selected BC markers were quantified. The effectiveness of pathway perturbation was quantified by expression of representative WNT, BMP and TGFb downstream targets. n = 3. (H) Representative flow cytometry plots showing the self-renewing capability of cells after the 5-day perturbation. EPCAM^+^/ITGB4^+^ double positive candidate eBCs (DP) and EPCAM^+^/ITGB4^-^ single positive cells (SP) were sorted and expanded on collagen peptide coated plates. After 3 passages, DP and SP groups maintained 83.2% and 15.4% candidate eBC cells, respectively. (I) IF staining of Calcium-induced stratification assay. The effect of Calcium treatment was evaluated by the observation of KRT13^+^/KRT4^+^ squamous cells, which are large and flat phenotypically. (J) Whole-mount IF staining of 3D organoids derived from DP cells showing KRT5^+^ eBCs and KRT13^+^ suprabasal cells. (K) IF staining of organotypic culture sections. hPSC-derived eBCs expanded and differentiated to esophageal squamous cells on decellularized dermis. Scale bars: 10 μm.

### The Manatee-predicted combination of local mesenchyme signals efficiently promotes eBC derivation from hPSCs

With the nominated *in vivo* signals as input, Manatee predicted that activation of EGF, BMP and TGFb along with inhibition of WNT (strategy #19) promote BC specification, which could enhance eBC derivation from hPSCs *in vitro*. In our modified hPSC differentiation system, although definitive endoderm (DE) and esophageal progenitors (EPC) could be efficiently derived, eBC efficiency dropped drastically at D43 (> 90% DE at D4, > 60% EPCAM^+^/P63^+^ EPC at D16, in contrast to ∼3% EPCAM^+^/ITGB4^+^ eBC at D43. Figure S7E). Such drop in eBC efficiency hindered manufacturing of esophageal mucosa at the clinical scale, mainly due to inefficient eBC specification from EPCs and fast expansion of non-epithelial cells over prolonged culture. To test whether strategy #19 could effectively and efficiently produce eBCs, we compared the D24 EPCs treated with EGF, BMP and TGFb ligands BMP4, TGFB1, and WNT antagonist IWP2 for another 5 days to those supplemented with EGF only (strategy #14 as the base control).

We also tested strategy #9 as the negative control, using EGF, BMP and TGFb inhibitors A83-01, DMH-1 and WNT agonist CHIR99021 (Figure 7A). Flow cytometry quantification confirmed that the Manatee-predicted strategy #19 indeed increased the proportion of ITGB4^+^/EPCAM^+^ eBCs in the EPCAM^+^ epithelial population compared to the base control #14 (38.63% ± 4.8% vs. 28.67% ± 1.56%). Notably, the negative control #9 drastically suppressed ITGB4^+^/EPCAM^+^ eBC formation (3.42% ± 0.56%. Figures 7B-C). Immunostaining showed only few cells expressing weak KRT5 by D29 (D24+5) in the #14 base control. In contrast, large quantities of strongly-staining KRT5^+^ cells were observed in the #19 treated D29 culture (Figure 7D), supporting the Manatee prioritization. No KRT5^+^ cells could be found in the #9 treated culture, further suggesting that activation of BMP/TGFb signaling and repression of WNT are required for the eBC commitment from EPCs. Consistently, #19 treated cells increased expression for the canonical BC markers KRT5 and KRT15 compared to #14 treated cells (Figures 7E-F). To comprehensively characterize the perturbation effects, we performed bulk RNA-seq for D24 culture, as well as the D29 cultures with 5-day treatment of #14, #9 and #19. Assessment of the corresponding signaling target transcripts confirmed effective activation of BMP/TGFb and repression of WNT by #19 and the reversed effect by #9. In support of the Manatee predictions, the expression of multiple BC marker genes, including TP63, ITGB4, ITGA6, COL7A1, COL17A1, KRT15 and KRT17, were all strongly increased in #19 treated cells, and largely repressed in #9 treated ones (Figure 7G).

Next, we tested whether #19-derived ITGB4^+^/EPCAM^+^ eBCs could function as their *in vivo* counterparts to self-renew and undergo squamous differentiation in culture and tissue organoid assays. We sorted the ITGB4^+^/EPCAM^+^ population (DP) from D29 culture and expanded the cells on collagen peptide coated plates for several passages. These expanded cells could maintain high expression of ITGB4, while the ITGB4^-^/EPCAM^+^ population (SP) remained largely absence of ITGB4 (Figure 7H), confirming the self-renewing capacity of #19-derived eBCs *in vitro*. After expansion, these ITGB4^+^/EPCAM^+^ eBCs generated esophageal-specific KRT13^+^ and KRT4^+^ suprabasal descendants when cultured over confluency (Figure 7I, -Ca^2+^). Notably, after Ca^2+^ treatment, large flattened KRT4^+^/KRT13^+^ could be observed that exhibited the typical squamous cell morphology. Hence, the #19-derived eBCs could respond to Calcium influx and undergo esophageal specific squamous stratification. To test whether ITGB4^+^/EPCAM^+^ eBC could generation 3D tissue organoids, we seeded the cells in Matrigel and cultured for 2 weeks. Remarkably, solid organoids were formed with KRT5^+^ BCs in the periphery and KRT13^+^ suprabasal cells inside (Figure 7J). These organoids could be passaged after dissociation into single cells, suggesting maintenance of self-renewing eBCs in the tissue organoids. In addition, we determined whether the eBCs could regenerate tissue-specific squamous stratified epithelium by growing ITGB4^+^/EPCAM^+^ eBCs on devitalized de-epidermal human dermis. Stratified skin basal cells will express KRT1 and KRT10, proteins needed to maintain skin integrity. Indeed, eBCs contributed to building the the COLVII^+^ basement membrane and generated KRT13^+^, KRT4^+^ and IVL^+^ squamous descendants on top of KRT5^+^/KRT15^+^/ITGB4^+^ basal layer. All of these cells were positive for human specific nuclei staining (huNuclei) as well as esophageal specific transcription factor PITX1, confirming their maintenance of tissue specificity even with the support of skin dermis (Figure 7K). Taken together, we successfully translated the *in vivo* human tissue interactive signals revealed from our atlas to the *in vitro* hPSC-to-eBC differentiation system using Manatee. The Manatee-prioritized strategy 19, activation of EGF, BMP, TGFb while inhibition of WNT, accelerated the derivation of functional eBCs and enhanced the efficiency, thus promising the manufacturing of esophageal mucosa at the clinical scale.

## DISCUSSION

Taking advantage of single cell and spatial technologies, we have created a robust multi-omics atlas of human tissue morphogenesis. Through integration with machine learning approaches, we have built an innovative framework to design actionable hPSC-directed differentiation strategies. Our multi-omics atlas systematically elucidates the developmental origin of eBCs and reveals two differentiation waves of epithelial morphogenesis accompanied by dynamic stromal architecture. By integrating spatial dynamics, we nominate local mesenchymal inductive signals driving eBC development as input for Manatee screening. Leveraging this framework, we establish the first human developmental signal-inspired tissue manufacturing system for esophageal mucosa. This promises the esophageal cell replacement therapy for RDEB as well as severe epithelial wounds caused by cancer resection or caustic injury. Our framework establishes a paradigm shift in the flexible design of human tissue manufacturing for clinical use.

Our atlas uncovers three unexpectedly complex yet orchestrated cellular dynamics during human esophageal development. First, given that our dataset has sampled developmental timepoints as early as the mid 1^st^ trimester, we for the first time clarified the molecular characteristics of the esophageal progenitors (Epi_PG) before the appearance of any differentiated cell types including eBCs. Particularly, we reveal GATA6 as an Epi_PG specific transcription factor. GATA6 plays essential roles in establishing endoderm in early development^30,31^ and later GATA6 expression becomes restricted posteriorly to gastric, intestinal and colonic epithelium but absent from the esophagus^79^. Notably, re-emergence of GATA6 expression has been found in Barrett’s esophagus and esophageal adenocarcinoma, characterized by the pathological conversion of stratified epithelium to intestinal metaplastic simple epithelium^80,81^. Our atlas could be leveraged in conjunction with disease and cancer databases to investigate any potential reactivation of developmental programs, and the role for the tumor microenvironment, during cancer progression.

Second, we reveal two differentiation waves in human esophageal epithelial morphogenesis, with ciliogenesis waning towards the end of 1^st^ trimester, and squamous stratification taking over afterwards. Ciliated esophageal epithelia are maintained in fish, amphibians and some reptiles, while their replacement by squamous stratified epithelia are observed in mammalian development^82^. We speculate that ciliated cells may help the fetus to swallow amniotic fluid, which is critical for the gastrointestinal development. Meanwhile, squamous cells become indispensable as the protective barrier for esophagus later. Interestingly, this transient appearance of ciliated cells in the esophagus is reminiscent of the periderm during skin development. With a strikingly similar pattern of two differentiation waves, upon specification from surface ectoderm progenitors, skin BCs first generate a layer of periderm before committing to squamous stratification. The periderm may serve as a “Teflon” coat to prevent pathological adhesion of adjacent epithelia before the establishment of the squamous cornified barrier^83,84^. These organ specific intermediates might reflect adaptation with distinct functional requirements, thus providing resources for evo-devo exploration.

Third, complex mesenchymal dynamics accompanies epithelial morphogenesis. Accompanying BC specification from Epi_PG, Fib_PG differentiates into 4 layers: Fib_1 being closest to the basement membrane, and MF, Fib_2, Fib_3 are further away, respectively. Notably, the drastic expansion of the Fib_2 layer in the submucosa distances the Fib_3, SM, and Fib_ICC containing muscularis externa, thus physically insulating morphogens secreted by muscularis externa from the epithelium. Additionally, it has been reported that DCN, a matrix component abundantly secreted by Fib_2, could antagonize SMAD signaling^85^. Hence, the mesenchymal cellular dynamics alter the tissue architecture. This results in stage-dependent changes in the cellular components of the local mesenchyme, which potentially drive BC development. While the local mesenchyme cell types remain the same from the mid to late stage, these cells could change their secreted morphogen profiles as they mature. Specifically, MF cells begin to express GREM2, an effective SMAD antagonist, from the late stage, rendering the epithelium in a SMAD inhibiting microenvironment to promote BC self-renewal^86,87^.

Facilitated by Manatee, we prioritized the *in vivo* local mesenchymal signals with ligands and small molecules to manufacture eBC *in vitro*. Instead of relying on prior knowledge gained using model animals, we here combined tissue atlas and machine learning to design clinical compatible hPSC differentiation system. The hPSC-to-eBC together with our previous hPSC-to-skin BC differentiation systems provide a unique chance to interrogate regulatory modules controlling the tissue-specific BC development. Our previous work revealed that during epidermal lineage commitment, the morphogen-induced expression of TFAP2C primes the chromatin landscape. Meanwhile, TFAP2C activates the canonical BC master regulator P63, which further effect on the primed chromatin for stratified epithelium maturation^88,89^. Our current work suggests a similar regulatory mechanism operates in endoderm. Specifically, induced key endodermal TFs such as SOX2 and GATA6 shape the endoderm-specific chromatin landscape as well as inducing P63; in turn, P63 induces stratified epithelium while repressing GATA6 to drive differentiation forward. Our functional assays placing esophageal basal cells on skin dermis provide strong evidence for the distinct and stable cellular specification network. Future mechanistic and evolutionary studies interrogating shared regulatory logic will help accelerate the production of other stratified epithelia such as bladder and cornea, as well as the mechanism of the master regulator P63 in specifying tissue-specific stratification programs in ectoderm and endoderm.

Our work highlights the broad applicability of Manatee to prioritize identified intercellular signals for tissue induction between tissue states. Besides screening for stem cell directed differentiation strategies, it could also be used to prioritize therapeutics to reverse diseased cellular states, or infer side effects of drugs. However, the Manatee neural network in its current architecture cannot appreciate TF-TF regulations. Therefore, during in silico perturbation, besides the candidate TF, all its downstream target TFs need to be adjusted for reliable predictions. We tackled this challenge by manually curating Gene Ontology (GO) pathway gene sets ^90, 91^ based on previous literature, thereby highlighting the need for more expansive and detailed TF-target databases on which to train the algorithm. In addition, next generations of Manatee will incorporate gene regulatory network topology in the design of the VAE architecture, similar to what has been done in the VEGA model^92^.

In summary, our multi-omics deepens our understanding in the development of human upper gastrointestinal system. The comprehensive characterization of cellular diversity and dynamics here provides a rich resource of *in vivo* landmarks for *in vitro* human cell engineering towards lineages across germ layers, including enteric neurons and glia cells, esophageal vascular components and fibroblasts. Instead of using ligands and small molecules, we could potentially co-derive or assemble the supporting cell types to further enhance manufacturing efficiency of esophageal mucosa. Moreover, our benchmarked atlas-spatial analysis-Manatee framework serves as a new paradigm in leveraging the human developmental cell atlas for the rational design of hPSC differentiation.

## METHODS

### Human Samples

Normal, de-identified human embryonic and fetal tissues (esophagus with stomach) were obtained from the NIH-funded University of Washington Birth Defect Research Laboratory (BDRL). All procedures were approved by the University of Washington and Stanford University Institutional Review Board (IRB#46721; SCRO#801). Tissue samples were shipped overnight in HBSS with ice packs. Detailed tissue sample information can be found in Table S1.

### Human Pluripotent Stem Cells

Human pluripotent stem cell (hPSC) line H9 was obtained from Stanford Stem Cell Bank. The RUES2 line (Rockefeller University Embryonic Stem Cell Line 2, NIH approval number NIHhESC-09-0013) was from Rockefeller University. All experiments using hPSC lines were approved by Stanford University. The H9 line was maintained in Essential 8 media (Thermo Fisher Scientific) on culture plates coated with iMatrix-511 (Takara). The RUES2 line was initially maintained on mouse embryonic fibroblasts (MEFs, irradiated CF-1 MEF, Thermo Fisher Scientific) in the maintenance medium: 400 ml of DMEM/F12 (Thermo Fisher Scientific), 100 ml of KnockOut Serum Replacement (Thermo Fisher Scientific), 5 ml of GlutaMAX (Thermo Fisher Scientific), 5 ml of MEM non-essential amino acids solution (Thermo Fisher Scientific), 3.5 ml of 2-mercaptoethanol (Sigma-Aldrich), 1 ml of primocin (Thermo Fisher Scientific), and FGF2 (R&D Systems) with a final concentration of 20 ng/ml to make a total of 500 ml of medium. For the feeder-free chemical-defined condition, the RUES2 cells were thawed and plated on culture plates coated with iMatrix-511 (Takara) in StemFit Basic04 Complete medium (Amsbio). For passaging, hPSCs were detached with TrypLE Select (Thermo Fisher Scientific) and plated at 1M per 10 cm dish. hPSCs were maintained in an incubator with 95% humidity, 95% air and 5% CO2 at 37 °C, and routinely tested for mycoplasma contamination.

### Immunocompromised Mice

Immunodeficient NIH-III Nude mice (Crl:NIH-Lyst^bg-J^Foxn1^nu^Btk^xid^) for skin patch transplantation and kidney capsule transplantation were purchased from Charles River (Strain Code 201). All mouse work was approved by the Institutional Animal Care and Use Committee (IACUC) at Stanford University.

### Human Esophageal Tissue Histology Staining

Human embryonic and fetal esophageal and stomach tissues were washed in cold HBSS. Whole esophageal tubes were cut into ∼3-5 mm length pieces from proximal to distal. Such processed tissues were subjected to both paraffin and OCT embedding, followed by H&E staining and imaging.

For paraffin embedding, tissues were fixed in 4% paraformaldehyde at 4 °C rotating overnight. Then tissues were washed in cold PBS for 3 times, transferred to 70% ethanol and sent to Human Pathology/Histology Service Core at Stanford for embedding, sectioning and H&E staining.

For OCT embedding, tissues were transferred to 30% sucrose in PBS at 4 °C rotating overnight till tissue sinking. Then tissues were transferred to 30% 1:1 sucrose-OCT solution at 4 °C overnight, and embedded in OCT the next day. During H&E staining, sections were initially warmed up to room temperature for 10 min, rehydrated in PBS for 10 min, and rinsed in water for 1 min. Then sections were stained in Hematoxylin (Millipore Sigma) for 30 sec, rinsed in water, and stained in Bluing reagent (Dako) for 30 sec, rinsed in water. Following that, sections were soaked in 50% ethanol for 1 min, 70% ethanol for 1 min, 95% ethanol for 1 min, Eosin (Millipore Sigma) for 1 min, 95% ethanol for 1 min twice, 100% ethanol for 1 min twice, Histo-Clear (Fisher Scientific) for 5 min, and fresh Histo-Clear overnight. On the following day, slides were mounted using Permount mounting medium (Fisher Scientific).

H&E images were acquired using AxioImager (Zeiss, Neuroscience Microscopy Service at Stanford).

### Immunofluorescence Staining and Microscopy Imaging

For immunofluorescence of coverslip cell cultures, cells were fixed in 4% paraformaldehyde at room temperature for 20 min and washed with PBS for 3 times. Cells were permeabilized and blocked with 0.3% Triton X-100 (Sigma-Aldrich) plus 10% normal horse serum (Jackson ImmunoResearch) in PBS for 1 hr. Primary antibodies were added into 0.3% Triton X-100 and 1% bovine serum albumin (Sigma-Aldrich) in PBS and incubated at 4 °C overnight. The next day, cells were washed with PBS and incubated with secondary antibodies and NucBlue Live ReadyProbes Reagent (Hoechst 33342, Thermo Fisher Scientific) at room temperature for 1 hr. After washing, cells on coverslips were mounted onto microscope slides using ProLong Gold Antifade Mountant. Antibody information is listed in Table S2.

For immunofluorescence of fixed frozen tissue sections, 7 μm frozen sections were warmed up to room temperature for 10 min and rehydrated in PBS for 10 min. Following that, sections were blocked and stained as described above.

Fluorescent images were taken using Leica Sp8 (Leica Microsystems, Cell Sciences Imaging Facility at Stanford) and LSM710 (Zeiss, Neuroscience Microscopy Service at Stanford) confocal laser scanning microscope.

### Chromium Single Cell Profiling

We performed Chromium single cell profiling on both primary samples and hPSC cell cultures. Specifically, human embryonic and fetal esophageal and stomach tissues were washed in cold HBSS. Whole esophageal tissues were collected and cleaned up by removing stomach tissues slightly proximal from the gastroesophageal junction and other connective tissues. For samples older than 8 post-conception weeks, esophageal tubes were cut open using fine scissors, and gently digested in 6 cm dishes with 25 U/ml Dispase (Corning) + 100 μg/ml DNase I (Millipore Sigma) at room temperature for 20 to 40 min. For samples younger than 8 post-conception weeks, whole esophageal tubes were digested in Dispase/DNase I solution at room temperature for 10 min instead. Digestion was stopped by transferring esophagus to cold HBSS. Epithelium was physically peeled off from surrounding stromal tissues with fine forceps, and collected in a 15 ml Falcon tube. The epithelium was further dissociated into single cells in TrypLE Express (Thermo Fisher Scientific), first triturated gently using P1000 pipette and then shaken in a Thermomixer at 37 °C at 1000 rpm for 10 to 20 min. Dissociation was stopped by addition of the FACS buffer (PBS + 1% BSA + 2 mM EDTA + 1X Anti-Anti + 25 mM HEPES pH 7.0) supplemented with 10% FBS. Stromal tissues were transferred to clean dishes and chopped into small pieces using scissors. Tissue pieces were transferred to a 15 ml Falcon tube with 10 ml collagenase solution (5 mg/ml Collagenase in DKSFM), triturated gently using P1000 pipette, and then shaken in a Thermomixer at 37 °C at 1000 rpm for 10 to 30 min. 1 ml 0.25% Trypsin was then added to the solution for an additional 5 min dissociation. Dissociation was stopped by addition of the FACS buffer supplemented with 10% FBS. Dissociated cells were resuspended in cold 0.04% BSA in PBS and filtered through 40 μm strainers. Cells were transferred to 1.5 ml Eppendorf tubes and pelleted down at 300 g at 4 °C for 5 min. Then cells were resuspended in cold 0.04% BSA in PBS for cell number counting. Cell alive rate was determined using Trypan Blue. Only samples with greater than 80% cell alive rate were further processed. Cells were then subjected to 10X Chromium scRNA-Seq based on the manual.

As for hPSC cell cultures, cells were dissociated using TrypLE Select, and resuspended in FACS buffer with SYTOX Blue Dead Cell Stain (Thermo Fisher Scientific). Live cells were sorted using FACSAria II (BD Biosciences) at Stanford Shared FACS Facility and collected in the FACS buffer supplemented with 10% fetal bovine serum (Thermo Fisher Scientific) and 2X Anti-Anti on ice. The sorted cells were pelleted, washed with cold 0.04% BSA in PBS, and resuspended in cold 0.04% BSA in PBS for cell number counting. Cell alive rate was determined using Trypan Blue. Only samples with greater than 90% cell alive rate were further processed. Cells were then subjected to 10X Chromium scRNA-Seq based on the manual.

Sequencing libraries were prepared using Chromium Single Cell 3’ v3 (CG000183 Rev C) and v3.1 (CG000315 Rev A) protocols according to the manufacturer’s manual. The pooled, 3’-end libraries were sequenced using Illumina NovaSeq. Cell Ranger version 4.0.0 was used for primary data analysis, including demultiplexing, alignment, mapping and UMI counting. Specifically, for alignment and mapping, the GRCh38 reference genome and corresponding annotation were used.

### scRNA-Seq Bioinformatics

Seurat version 4.0.3 was used for single cell gene expression quantification, dimension reduction, clustering analysis and marker gene identification ^93^ . Monocle version 2.18.0 was used for single cell trajectory analysis^94^. In order to perform efficient monocle analysis, without losing generality, we limit the maximum number of single cells per cell type as 1000. CellChat was used for single cell intercellular communication analysis^76^. Same as the monocle analysis, we limit the maximum number of single cells per cell type to 1000. Seurat, monocle and CellChat analyses were performed under R version 4.0.2.

### Visium Spatial Transcriptomic Profiling

Human embryonic and fetal esophageal and stomach tissues were washed in cold HBSS. Whole esophageal tubes were cut into ∼3-5 mm length pieces from proximal to distal. Fresh tissues were directly embedded in OCT on dry ice. 10 μm fresh frozen sections of OCT-embedded tissues were used for Visium profiling. Such sections were collected using cryostat, sealed in airtight 50 ml Falcon tubes, stored at -80 °C for downstream profiling.

Sequencing libraries were prepared using the Visium Spatial 3’ v1 (CG000239_Rev F) protocol according to the manufacturer’s manual. The pooled, 3’-end libraries were sequenced using Illumina NovaSeq. Space Ranger version 1.3.1 was used for primary data analysis, including demultiplexing, alignment, mapping and UMI counting. Specifically, for alignment and mapping, the GRCh38 reference genome and corresponding annotation were used.

### Cell2location Spatial Mapping Analysis

The cell2location algorithm maps all identified single cell populations as listed in Figure 3A onto Visium spatial transcriptomic profiles^72^. In order to make accurate spatial mapping, we 1) merged several groups of closely-related cell types, and 2) analyzed early/mid stage and late stage profiles separately. In order to make efficient spatial mapping, without losing generality, we limit the maximum number of mapped single cells per cell type as 500. The cell2location analysis was performed under Python version 3.9.12.

### CODEX Multiplexed Immunofluorescence Staining

CODEX profiling starts with antibody conjugation, which was performed using protocols and reagents per manufacturer instructions (Akoya; 7000009). Such a process conjugates 50 μg of immunofluorescence-validated carrier-free antibodies to specific barcodes (see Table S3). Specifically, antibodies were concentrated on a 50 kDa filter equilibrated with the filtration buffer. The sulfhydryl groups were activated by incubating for 30 mins at room temperature with the reduction mix. Antibodies were then washed with the conjugation buffer once. Oligonucleotide barcodes were resuspended in the conjugation buffer, added to the antibodies, and allowed to incubate for 2 hrs at room temperature. The conjugated antibodies were washed 3 times, by resuspending and spinning down at 12,000 g for 8 mins with the purification solution. Antibodies were then eluted by adding 100 μl storage buffer and spinning at 3000g for 4 mins. The conjugated antibodies were stored at 4°C till use.

Before use in multiplexed CODEX experiments, conjugated antibodies were validated with CODEX single stains on human fixed frozen fetal esophageal tissues. Staining was performed with the conjugated antibody as described below. The screening buffer was prepared according to the CODEX manual provided by Akoya Biosciences. Fixed and stained tissues were incubated in the screening buffer for up to 15 mins. Fluorescent DNA probes were prepared and added to stained tissues for 5 mins. Tissues were washed 3 times with the screening buffer followed by 1 wash with the CODEX buffer. Tissues were then imaged using a Keyence BZ-X810 inverted fluorescent microscope for the antibody validation.

All fixed frozen tissue CODEX antibody stainings were done according to Akoya Biosciences staining protocol associated with the CODEX Staining Kit with some modifications (Akoya; 7000008). After tissue hydration, autofluorescence was bleached using a previously published protocol by allowing the tissue to sit in a solution (made of sodium hydroxide and hydrogen peroxide) sandwiched between a LED light panel for 45 mins at room temperature^95^. The tissue was then washed twice in ddH2O for 10 mins, followed by a wash in the CODEX Hydration Buffer (contained in the CODEX Staining Kit) twice, 2 mins each. The coverslip was allowed to equilibrate in the CODEX staining Buffer (contained in CODEX Staining Kit) for 20 mins at room temperature. The blocking buffer was prepared by adding N, S, J, and G blocking solutions to the CODEX staining buffer (contained in the CODEX Staining Kit). Antibodies were added to the blocking buffer to make a total volume of 200 μl. The antibody cocktail was added to the coverslip, and staining was performed in a sealed humidity chamber at 4°C overnight. The antibody clones and titrations used are listed in Table S3. After staining, coverslips were washed twice in the hydration buffer for 4 mins followed by fixation in the storage buffer (contained in the CODEX Staining Kit) with 1.6% paraformaldehyde for 10 mins. Coverslips were then washed thrice in PBS, followed by a 5 min incubation in ice-cold methanol on ice for 5 mins followed by another 3 1X PBS washes. CODEX fixative solution (contained in CODEX Staining Kit) was prepared right before the final fixation step. 20 μl of CODEX fixative reagent was added to 1000 μl 1X PBS. 200 μl fixative solution was added to the coverslip for 20 mins followed by 3 washes in 1X PBS. Coverslips were then immediately prepped for imaging.

For CODEX multicycle setup and imaging, coverslips were mounted onto Akoya’s custom-made stage between coverslip gaskets with the tissue side facing up. The coverslips were cleaned from the bottom using a Kim wipe to get rid of any salts. The tissue was stained with Hoechst Nuclear Stain (Thermo Fisher Scientific, cat. no. 62249) at a 1:2000 dilution in the 1X CODEX buffer. A 96-well plate was used to set up the multicycle experiment with different fluorescent oligonucleotides in each well. A reporter stock solution was prepared to contain 1:2000 Hoechst stain and 1:12 dilution of assay reagent in the 1X CODEX buffer (contained in the CODEX Staining Kit). Fluorescent oligonucleotides (Akoya Biosciences) were added to this reporter solution at a final concentration of 1:50 in a total of 250 μl per well. A blank cycle containing no fluorescent probes was performed at the start and end of the experiment to capture residual autofluorescence (see Table S3 for the cycle set up). Automated image acquisition was performed using the CODEX Instrument Manager (CIM, version 1.30, Akoya Biosciences). Imaging was performed using a Keyence BZ-X810 microscope, fitted with a Nikon CFI Plan Apo l 20X/0.75 objective. The BZ-X software (Keyence) multi-point option was used to define the center and the imaging area corresponding to each region. 9-11 z steps were acquired with the pitch set at 1.5 in the BZ-X software. Processed images were outputted in tiff format for downstream analysis.

### CODEX Analysis

Raw tiff files were processed using the CODEX Processor version 1.7.0.6 by Akoya Biosciences. CODEX Processor sequentially performs drift-compensation, deconvolution, background subtraction, tile stitching and segmentation. Such a pipeline yields spatial coordinates as well as marker protein signal intensities for all the detected cells. We then performed quality control based on the average signal intensity of all the DAPI channels. Cells with low average DAPI signals were removed, and the remaining cells were subjected to signal intensity log-transformation, hierarchical clustering and marker-guided annotation.

### Manatee

The Manatee deep learning framework was adapted from VAE^77^ and designed to model the generative process from TF expression to the corresponding gene expression. In order to do so, 1) both the encoder and decoder neural networks consist of two fully connected layers, each with the same number of nodes as the number of genes; 2) the latent space contain the same number of variables as the number of TFs; and 3) the following loss function is optimized during training:

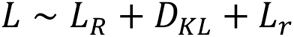

where *LR* and *DKL* represent reconstruction loss and Kullback–Leibler Divergence against the *N(**0**, **1**)* normal distribution respectively, as the two regular VAE loss terms. The additional *Lr* term represents the TF reconstruction loss, which is the mean square error (MSE) between reparameterized latent variables (***Z***) and TF expression (***TF***):

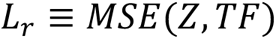

Manatee analysis starts with the TF expression matrix (D16 and D24 in our case), and perturbations are made by adjusting values of corresponding latent variables. The adjusted TF expression profiles are then passed through the decoder to predict expression profiles after perturbations. In order to assign biologically legitimate values to candidate latent variables, values will be sampled from the reference matrix, which tracks the TF expression pattern of the target phenotype (BC in our case). We performed a TF-wise sampling of the reference matrix, which gets values from only the corresponding TF expression vector. We also specified the perturbation direction and the sampling quantile for each TF. For up-regulated TFs, expression values will be sampled from the top quantile, and *vice versa* for down-regulated TFs.

We defined TFs based on the Gene Ontology (GO) database^90,91^, under terms GO:0003700, “DNA-binding transcription factor activity”, GO:0003677, “DNA binding”, GO:0140110, “transcription regulator activity” and GO:0006355, “regulation of DNA-templated transcription”. The full human TF list is available as Table S4.

We also downloaded pathway gene lists from the GO database, including terms GO:0007173, “epidermal growth factor receptor signaling pathway”, GO:0007179, “transforming growth factor beta receptor signaling pathway”, GO:0016055, “Wnt signaling pathway” and GO:0030509, “BMP signaling pathway”. When perturbing specific signaling pathways, the corresponding lists will be retrieved, and non-TFs will be filtered. The yielded lists will then be manually annotated based on previous literature in terms of determining perturbation directions (Table S5).

We trained Manatee by merging *in vivo* (D4, 10, 16, 24 and 43) and *in vitro* (Epi_PG, BC and SB-1 to 5) single cell datasets. We designed such a training to comprehensively capture regulatory logics during the BC specification and differentiation *in vivo*, as well as among the hPSC culture milestones *in vitro*. For each cell population, we downsampled the expression matrix to a maximum 1,000 cells. We performed 5 independent trainings and the one with the minimum loss value was used for downstream predictions.

### *In vitro* Differentiation of hPSCs into Esophageal Basal Cells

We adapted three previous protocols to differentiate hPSCs into esophageal progenitor cells in feeder-free chemically-defined conditions^18,19,78^. 0.1 million H9 or RUES2 hPSCs were plated in E8 medium with 10 μM Rock inhibitor Y-27632 (Tocris) in 1 well of 12-well culture plates coated with iMatrix-511. 24 hrs later, culture medium was changed to MCDB131 base medium supplemented with 5 μM CHIR99021 (Tocris) and 100 ng/ml Activin A (R&D System) for 24 hrs. MCDB131 base medium was prepared as following: 500 ml MCDB131 (Thermo Fisher Scientific), 5 ml of GlutaMAX (Thermo Fisher Scientific), 33 ml of 7.5% Bovine Albumin Fraction V Solution (Thermo Fisher Scientific), 10 ml of 7.5% NaHCO3 (Thermo Fisher Scientific) and 2 ml of 45% Glucose-D (Sigma Aldrich). On day 2, the medium was supplemented with 0.5 μM CHIR99021 and 100 ng/ml Activin A. On day 3, the medium was supplemented with 100 ng/ml Activin A only. On day 4, base medium was changed to Serum-Free Differentiation (SFD) medium: 750 ml of reconstituted IMDM (Thermo Fisher Scientific), 250 ml of F-12 (Corning), 7.5 ml of 7.5% Bovine Albumin Fraction V Solution (Thermo Fisher Scientific), 10 ml of GlutaMAX (Thermo Fisher Scientific), 5 ml of N2 (Thermo Fisher Scientific), 10 ml of B27 (Thermo Fisher Scientific) and 10 ml of Penicillin/Streptomycin (Thermo Fisher Scientific), and adding L-Ascorbic acid (Sigma-Aldrich) and MTG (Sigma-Aldrich) on the day of use to obtain a final concentration of 50 mg/ml and 0.04 ml/ml, respectively. Additional 1X GlutaMAX was also added. On day 4 and 5, cells were induced for anterior foregut endoderm differentiation in the SFD medium plus 10 mM SB431542 (Tocris) and 100 ng/ml Noggin (R&D Systems). From day 6 to day 9, anterior foregut endoderm was induced to dorsalize in the SFD medium supplemented with 10 mM SB431542, 100 ng/ml Noggin, 100 ng/ml hEGF (R&D Systems) and 50 ng/ml FGF10 (R&D Systems). From day 10 to day 15, esophageal progenitor cells were induced in the SFD medium supplemented with 100 ng/ml hEGF and 50 ng/ml FGF10. From day 16 onward, cells were cultured in the SFD medium supplemented with 100 ng/ml hEGF only.

To test pathway perturbation strategies predicted by Manatee, different combinations of growth factors and chemicals were added to the SFD medium with 100 ng/ml hEGF. Cells were treated in the perturbation conditions for 5 additional days from day 16 or 24. In experimental strategy #19, cells were cultured in the SFD medium supplemented with 100 ng/ml hEGF, 5 ng/ml BMP4 (R&D Systems), 2 ng/ml TGFB1 (R&D Systems) and 0.4 μM IWP2 (Selleck Chemicals). In negative control strategy #9, cells were cultured in the SFD medium supplemented with 100 ng/ml hEGF, 1 μM A83-01 (Tocris), 1 μM DMH-1 (Tocris) and 3 μM CHIR99021 (Tocris). In base control strategy #14, cells were cultured in the SFD medium supplemented with 100 ng/ml hEGF only.

### Flow Cytometry and Cell Sorting

For cell surface marker staining, cells were dissociated with TrypLE Select and stained with fluorophore conjugated antibodies in the FACS buffer for 20 to 30 min at room temperature. After washing in the FACS buffer, stained cells were resuspended in FACS buffer with SYTOX Blue Dead Cell Stain to exclude dead cells.

For intracellular staining, cells were processed using eBioscience Foxp3/Transcription Factor Staining Buffer Set (Thermo Fisher Scientific) according to the manufacturer’s instructions. Briefly, cells were stained with surface markers and Fixable Viability Dye eFluor 450 for 30 to 45 min at 4 °C. Fixation and permeabilization was performed at room temperature for 1 hr, followed by primary antibody incubation overnight at 4 °C, and secondary antibody incubation at room temperature for 1 hr.

UltraComp eBeads Compensation Beads were used for single color compensation control. Flow cytometric analysis was performed using LSR II (BD Biosciences) and cell sorting was performed using FACSAria II at Stanford Shared FACS Facility. Data was analyzed using FlowJo software. Cells were sorted into the FACS buffer supplemented with 10% fetal bovine serum and 2X Anti-Anti.

### qPCR

Cells were lysed with TRIzol Reagent (Thermo Fisher Scientific) and RNA was extracted using Direct-zol RNA Miniprep Plus kit (Zymo Research). Quantitative PCR was performed using the 1-Step Brilliant II SYBR Green QRT-PCR Master Mix kit (Agilent Technologies) and the TaqPath 1-Step Multiplex Master Mix kit (Thermo Fisher Scientific) using a LightCycler 480 instrument (Roche). Gene expression was normalized to the internal control β-actin or GAPDH. All qPCR experiments were performed with at least triplicates. KRT5 primers are AGAGCTGAGAAACATGCAGG (forward) and AGCTCCACCTTGTTCATGTAG (reverse); KRT5 primers are AGGACTGACCTGGAGATGCAGA (forward) and TGCGTCCATCTCCACATTGACC (reverse).

### Bulk RNA-Seq

RNA concentration and quality was measured by 2100 Bioanalyzer (Agilent Technologies). Libraries were prepared using SMART-Seq v4 Ultra Low Input RNA kit (Takara Bio) and sequenced by NovaSeq (Illumina) with paired-end 100 bp reads at MedGenome. Each library was sequenced with a targeted depth of 40 million total reads. 2 or 3 biological replicates were sequenced for each sample. Alignment was performed using STAR^96^ against the hg38 reference genome. RPKM gene expression values were quantified using HOMER analyzeRepeats.pl (http://homer.ucsd.edu/homer/ngs/analyzeRNA.html).

### Calcium Induced Stratification

Sorted EPCAM^+^/CD104^HI^ DP cells were expanded in Defined Keratinocyte Serum-Free Medium (DKSFM, Thermo Fisher Scientific) supplemented with 10 μM Y-27632, 1 μM A83-01 (Tocris) and 1 μM DMH-1 (Tocris) on 6-well ECM collagen-I-coated peptide plates (Corning) or culture plates coated with rat tail collagen-I solution (Cell Biologics). After confluency, cells were treated with 2 mM CaCl2 for 8 days.

### 3D Organotypic Culture

Devitalized de-epidermal human dermis (DED) was prepared as follows. Cadaver skin (New York Firefighter Skin Bank) was freeze–thawed three times to devitalize cells and washed in PBS with 5X penicillin-streptomycin, 5X gentamicin and 5X fungizone. The sterilized skin was stored in PBS containing 1X penicillin-streptomycin, 1X gentamicin and 1X fungizone at 37 °C for one week. The epidermis was then peeled off of the dermis, which was then stored in PBS containing 1X penicillin-streptomycin at 4 °C for more than two weeks before use. Devitalized dermis was cut into 1.5 cm × 1.5 cm pieces, and stored epidermis-down in 6-well dishes at 37 °C to let the dermis attach to the bottom. The culture was switched to DKSFM, and 10^6^ induced esophageal BCs were seeded onto the center of DED. After 3 days, the culture was lifted to the air–liquid interface and switched to KGM medium (DMEM: Hams F12 3:1, FBS 10%, 1X nonessential amino-acid, 0.18 mM adenine hydrochloride, 0.1 nM cholera toxin, 10 ng/ml EGF, 0.4 μg/ml hydrocortisone, 5 μg/ml insulin, 2 nM triiodo-l-thyronine, 5 μg/ml transferrin). After two to four weeks, the organotypic co-cultures were collected and the pieces were fixed in 4% PFA at room temperature for 4 hrs or 4 °C overnight, washed with PBS, and processed for OCT and paraffin embedding for downstream analysis.

### 3D Organoid Culture

60,000 EPCAM^+^/CD104^HI^ sorted DP cells were resuspended in 60 μl SFD medium supplemented with 100 ng/ml hEGF, 20 ng/ml FGF2, 10 μM Y-27632, 1 μM A83-01 (Tocris) and 1 μM DMH-1 (Tocris) and mixed with 90 μl Growth Factor Reduced Matrigel (Corning). The mixture was added in 24-well cell culture inserts (Falcon), and incubated at 37 °C incubator for 30 min to solidify. The medium was added to both the bottom and top chambers, and changed every 2 to 3 days.

### Data Availability

Bulk sequencing, scRNA-Seq profiles and Visium profiles are deposited on the dbGAP server under accession number phs003281.v1.p1. Raw CODEX images are available upon request to corresponding authors.

### Code Availability

The Manatee algorithm is available at https://github.com/hd2326/Manatee.

Customized scripts are available upon request to corresponding authors.

### Graphics

The diagrams in Figures 1A, 4C, 5A were created by the authors using Biorender.com.

## ACKNOWLEDGEMENTS

We thank the Oro lab for helpful discussions, Stanford Research Computing Facility (SRCF), the University of Arizona High Performance Computing team and the College of Pharmacy Information Technology Group for computational support. We thank Pauline Chu, BS, HT (ASCP), Stanford Human Histology Research Core, Stanford Genomics Core (supported by NIH S10OD025212 and 1S10OD021763) Stanford Neuroscience Microscopy Service, Wesley Alejandro and Anum Khan at the Stanford Cell Sciences Imaging Facility (RRID: SCR_017787), Stanford Shared FACS Facility (RRID: SCR_017788). The project described was supported, in part, by Award Number 1S10OD010580-01A1 from the National Center for Research Resources (NCRR). Its contents are solely the responsibility of the authors and do not necessarily represent the official views of the NCRR or the National Institutes of Health. We also thank the University of Washington the Birth Defects Research Laboratory for sample collection. A.E.O. is supported by NIAMS R01ARO73170, Department of Defense PR212394, EB Research Partnership, and Stanford Innovative Medicine Accelerator. H.D. is supported by the University of Arizona Health Sciences Career Development Award. I.A.G. is supported by NICHD R24HD000836. J.Q. is supported by DK132251, CA272901, HL152293.

## AUTHOR CONTRIBUTIONS

Y.Y. performed single cell RNA-Seq and Visium experiments. Y.Y., N.Y.L., performed CODEX experiments. Y.Y., Y.Z., J.Q. developed the standard hPSC differentiation protocol. Y.Y., C.G.M., W.J.K., J.T. performed hPSC to eBC differentiation experiments. Y.Y., C.P., H.Z. performed 3D organotypic culture. Y.Y., L.S., J.S. and H.D. developed the Manatee algorithm and performed the analysis. Y.Y., N.Y.L., S.G. and H.D. performed single cell, Visium and CODEX analyses. I.A.G. and the BDRL collected human samples. G.C. helped with pathology analysis. L.G. and H.D. built the data portal. H.D. and A.O. supervised the project. Y.Y., H.D. and A.O. wrote the manuscript. All authors read and approved the manuscript.

