## Supplementary material for "A Spatiotemporal and Machine-Learning Platform Accelerates the Manufacturing of hPSC-derived Esophageal Mucosa": 7 Supplementary Figures

#### Supplemental Data Items

**Figure S1. scRNA-seq data quality control and filtering, related to Figure 2.**
(A) For each dataset, sample-wise distributions of number of cells/spots, number of UMIs per cell and number of genes per cell were reported. (B) Iterative clustering of epithelium (EPI) and stromal (STROMA) single cell collections processed using TrpLE and Collagenase, respectively. (C-D) UMAP visualization of the coarse-grain initial clustering of EPI (C) and STROMA (D), color-coded based on cell type (left panel) and developmental timepoint (middle panel). Right panel: dot plots showing the expression of selected differentially expressed genes for each cellular compartment.

**Figure S2. Cellular heterogeneity in EPI and MES, related to Figures 2 and 3.**
(A) H&E staining of developing human esophageal epithelium at high resolution from E45 to E130. (B-C) 3 dimensional UMAP visualization of epithelial cells color-coded based on cell type/state (B) and developmental timepoints (C). (D) Dot plot showing the expression of epithelial differential marker genes. (E-F) 3 dimensional UMAP visualization of mesenchymal cells color-coded based on cell type (E) and developmental timepoint (F). (G) Dot plot showing the expression of mesenchymal differential marker genes.

**Figure S3. Cellular heterogeneity and lineage analysis of other stromal compartments, related to Figure 3.**
(A, C) 3 dimensional UMAP visualization of enteric nervous system (ENS) cells color-coded based on cell type/state (A) and developmental timepoint (C). (B) Proportions of the identified cell types at each timepoint. (D) Dot plot showing the expression of ENS differential marker genes. (E-F) Representative CODEX images showing the presence of ENS_PG at E45 (Early, E) and Neu and Glia at E120 (Late, F). (G) Monocle2 trajectory of the whole ENS compartment, color-coded based on cell type. (H, J) 3 dimensional UMAP visualization of endothelial (ENDO) cells color-coded based on cell type/state (H) and developmental timepoint (J). (I) Proportions of the identified cell types at each timepoint. (K) Dot plot showing the expression of ENDO differential marker genes. (L) Dot plot showing selected markers of Art, Ven, Lym, Peri and Im shown in CODEX images. (M-N) Representative CODEX images showing the presence of Lym, Art, Ven, Peri (M) and Im (N) at E120 (Late). Scale bars: 10 μm.

**Figure S4. CODEX antibody panel and cell type annotation reference, related to Figures 2-4.**
(A) Dot plot showing RNA expression of CODEX marker genes in different cell types. (B) Dendrogram workflow for cell type annotation using selected markers.

**Figure S5. Integrated analysis of single cell and Visium spatial transcriptomics, related to Figure 4.**
(A, C) Visium spatial feature plots visualizing selected cell types in different combinations on E72 (Mid, A) and E120 (Late, C) esophageal sections, color-coded based on mapping scores. (B, D) Visium spatial feature plots visualizing single cell type on E72 (Mid, B) and E120 (Late, D) esophageal sections, color-coded based on mapping scores.

**Figure S6. eBC driving signal combination screening by Manatee, related to Figure 6.**
(A) Quality control plot of *in vitro* single cell collections. (B-C) UMAP visualization of *in vitro* single cell collections color-coded based on cell type (B) and stage (C). (D) Original single cell expression profiles and predictions yielded by all 81 possible strategies were visualized on the two-dimensional UMAP space. Cells were color-coded by their types, and prediction effects were noted by arrows.

**Figure S7. Characterization of the chemically-defined, xeno-free hPSC-to-eBC differentiation system, related to Figure 7.**
(A) Principal-component analysis (PCA) of bulk RNA-seq data from hPSC-to-eBC differentiation at different timepoints. (B) Heatmap of differential gene expression along hPSC-to-eBC differentiation. Selected markers were labeled to the left. Right panel showing the top enriched ARCHS4_TISSUES terms by EnrichR. (C) IF of human esophageal transcription factors on D16 cells. (B) IF of BC markers and squamous markers on D24, D35, and D45 cells. Note that BC keratins are gradually acquired over the time course. (C) Representative flow cytometry quantification of differentiation efficiency, showing high efficiency in deriving DE on D4 and EPC on D16, but low efficiency in $EPCAM^{+}/ITGB4^{+}$ eBCs on D43. Scale bars, 10 µm.

**Table S1. Metadata information for human samples, related to Figure 1.**

**Table S2. Immunofluorescence staining antibody spreadsheet, related to Figure 7 and S7.**

**Table S3. CODEX configuration spreadsheet, related to Figure 4.**

**Table S4. Human TF list, related to Figure 6.**

**Table S5. Pathway TF list, related to Figure 6.**

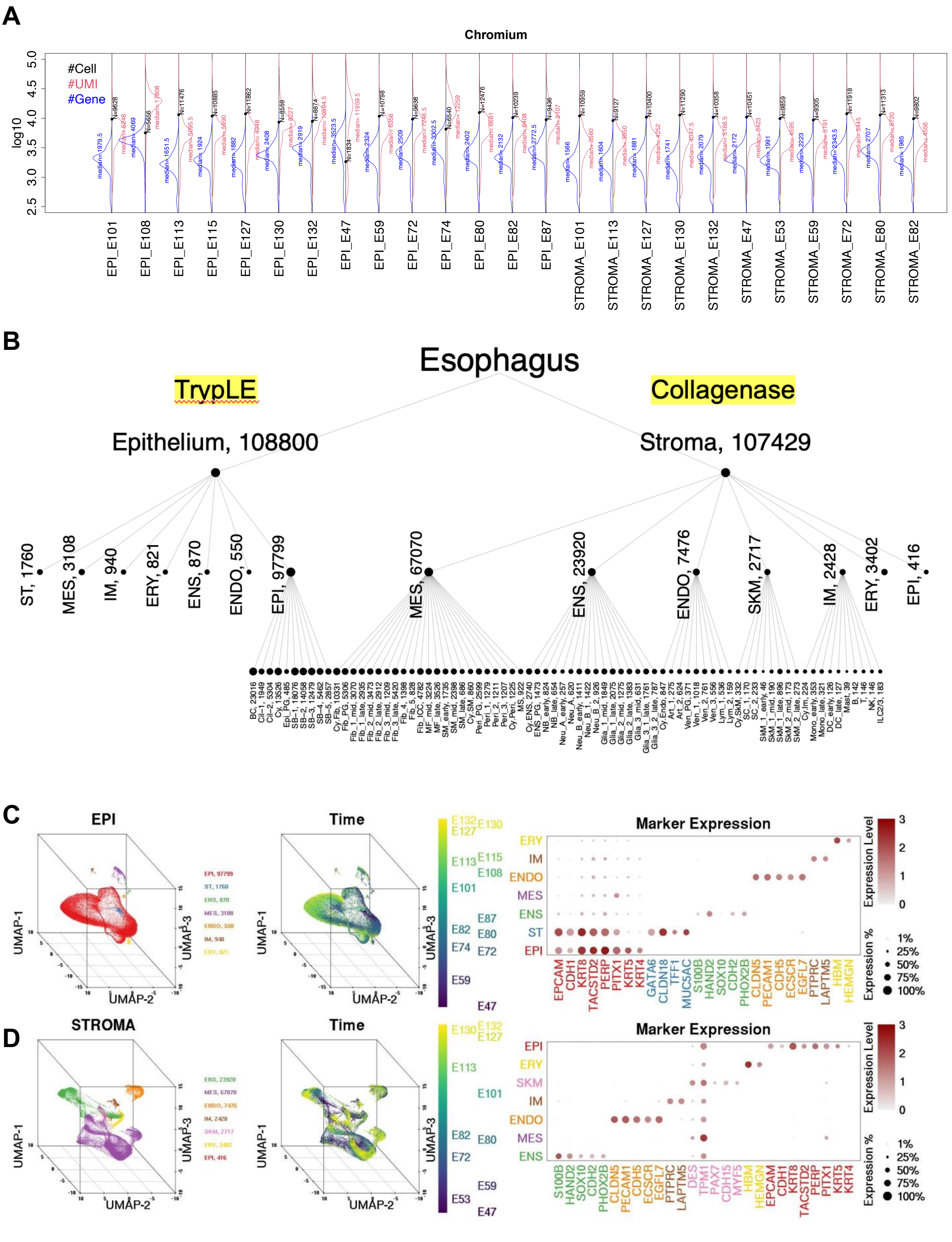
A
Chromium
#Cell
#UMI
#Gene
log10
EPI_E101
EPI_E108
EPI_E113
EPI_E115
EPI_E127
EPI_E130
EPI_E132
EPI_E47
EPI_E59
EPI_E72
EPI_E74
EPI_E80
EPI_E82
EPI_E87
STROMA_E101
STROMA_E113
STROMA_E127
STROMA_E130
STROMA_E132
STROMA_E47
STROMA_E53
STROMA_E59
STROMA_E72
STROMA_E80
STROMA_E82
B
Esophagus
TrypLE
Collagenase
Epithelium, 108800
Stroma, 107429
ST, 1760
MES, 3108
IM, 940
ERY, 821
ENS, 870
ENDO, 550
EPI, 97799
MES, 67070
ENS, 23920
ENDO, 7476
SKM, 2717
IM, 2428
ERY, 3402
EPI, 416
C
EPI
EPI, 97799
ST, 1760
ENS, 870
MES, 3108
ENDO, 550
IM, 940
ERY, 821
UMAP-1
UMAP-2
UMAP-3
Time
Marker Expression
ERY
IM
ENDO
MES
ENS
ST
EPI
EPCAM
CDH1
KRT8
TACSTD2
PERP
PITX1
KRT5
KRT4
GATA6
CLDN18
TFF1
MUC5AC
S100B
HAND2
SOX10
CDH2
PHOX2B
CLDN5
PECAM1
CDH5
ECSCR
EGFL7
PTPRC
LAPTM5
HBM
HEMGN
Expression Level
Expression %
1%
25%
50%
75%
100%
D
STROMA
ENS, 23920
MES, 67070
ENDO, 7476
IM, 2428
SKM, 2717
ERY, 3402
EPI, 416
UMAP-1
UMAP-2
UMAP-3
Time
Marker Expression
EPI
ERY
SKM
IM
ENDO
MES
ENS
S100B
HAND2
SOX10
CDH2
PHOX2B
CLDN5
PECAM1
CDH5
ECSCR
EGFL7
PTPRC
LAPTM5
DES
TPM1
PAX7
CDH15
MYF5
HBM
HEMGN
EPCAM
CDH1
KRT8
TACSTD2
PERP
PITX1
KRT5
KRT4
Expression Level
Expression %
1%
25%
50%
75%
100%

Figure S1

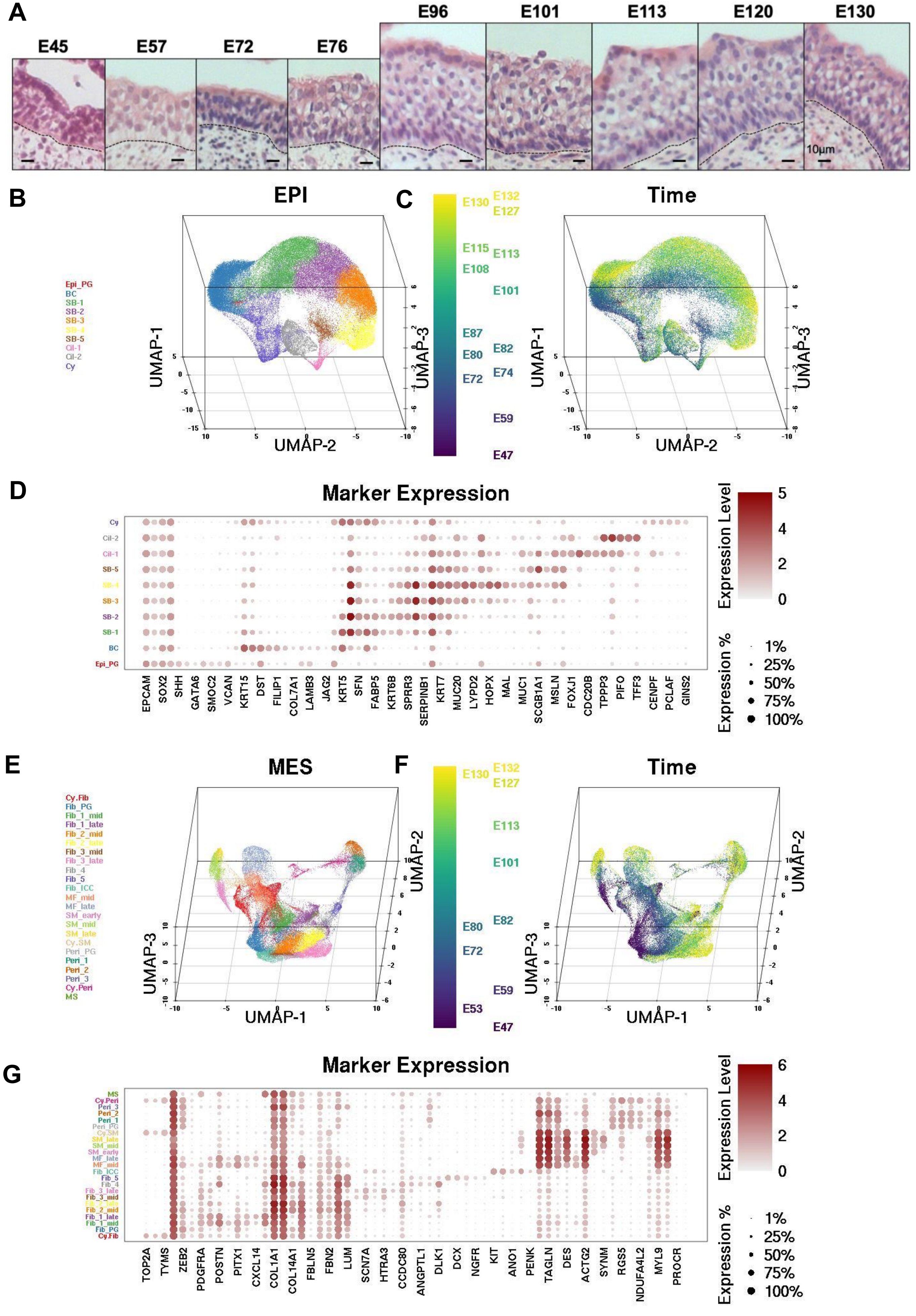
A
E45
E57
E72
E76
E96
E101
E113
E120
E130
10μm
B
EPI
Epi_PG
BC
SB-1
SB-2
SB-3
SB-4
SB-5
Cil-1
Cil-2
Cy
UMAP-1
UMAP-2
UMAP-3
C
Time
E130
E132
E127
E115
E113
E108
E101
E87
E82
E80
E74
E72
E59
E47
D
Marker Expression
EPCAM
SOX2
SHH
GATA6
SMOC2
VCAN
KRT15
DST
FILIP1
COL7A1
LAMB3
JAG2
KRT5
SFN
FABP5
KRT6B
SPRR3
SERPINB1
KRT7
MUC20
LYPD2
HOPX
MAL
MUC1
SCGB1A1
MSLN
FOXJ1
CDC20B
TPPP3
PIFO
TFF3
CENPF
PCLAF
GINS2
Expression Level
Expression %
1%
25%
50%
75%
100%
E
MES
Cy.Fib
Fib_PG
Fib_1_mid
Fib_1_late
Fib_2_mid
Fib_2_late
Fib_3_mid
Fib_3_late
Fib_4
Fib_5
Fib_ICC
MF_mid
MF_late
SM_early
SM_mid
SM_late
Cy.SM
Peri_PG
Peri_1
Peri_2
Peri_3
Cy.Peri
MS
F
Time
E53
G
Marker Expression
TOP2A
TYMS
ZEB2
PDGFRA
POSTN
PITX1
CXCL14
COL1A1
COL14A1
FBLN5
FBN2
LUM
SCN7A
HTRA3
CCDC80
ANGPTL1
DLK1
DCX
NGFR
KIT
ANO1
PENK
TAGLN
DES
ACTG2
SYNM
RGS5
NDUFA4L2
MYL9
PROCR

### Figure S2

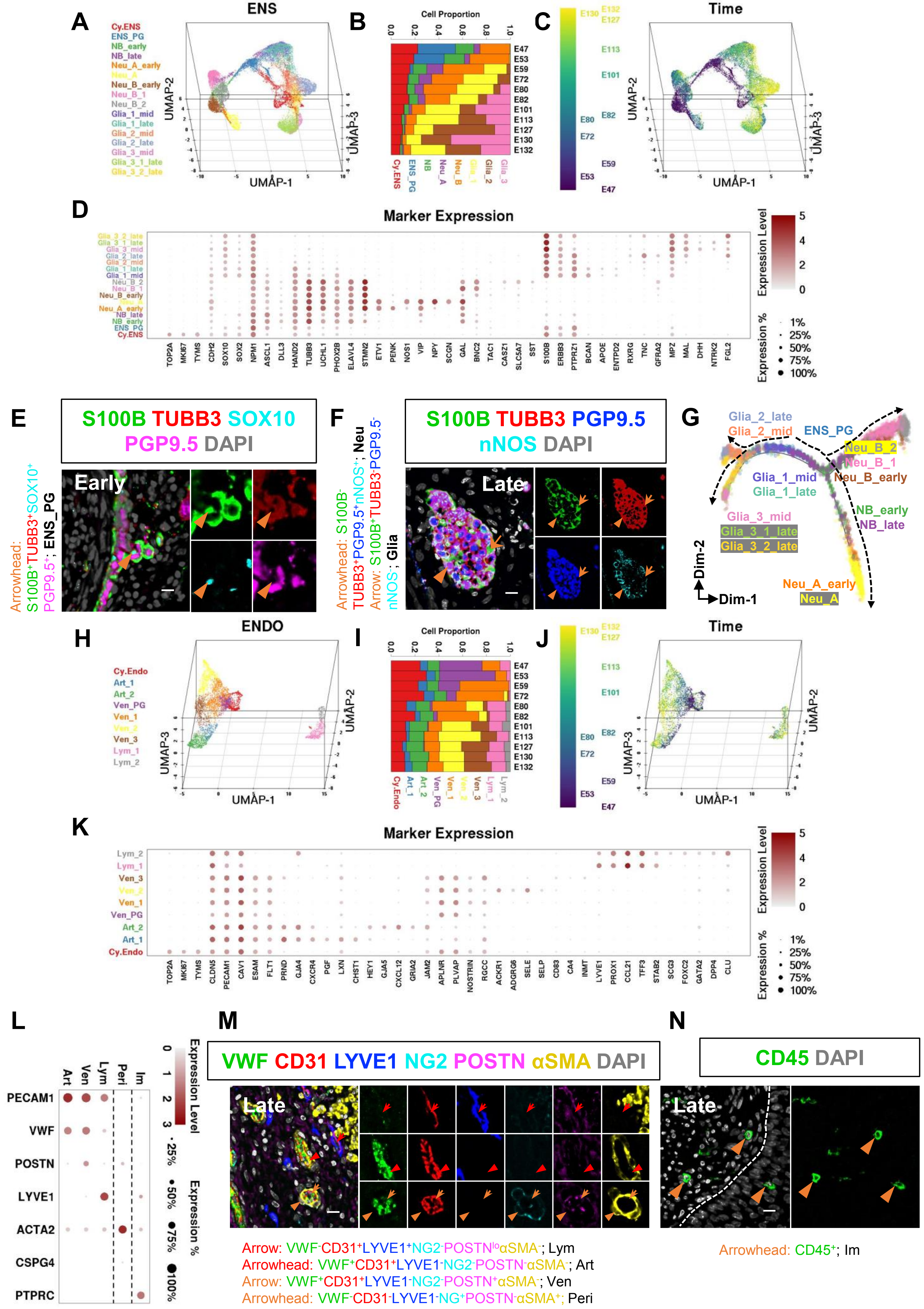
A
ENS
Cy.ENS
ENS_PG
NB_early
NB_late
Neu_A_early
Neu_A
Neu_B_early
Neu_B_1
Neu_B_2
Glia_1_mid
Glia_1_late
Glia_2_mid
Glia_2_late
Glia_3_mid
Glia_3_1_late
Glia_3_2_late
UMAP-1
UMAP-2
UMAP-3
B
Cell Proportion
E47 E53 E59 E72 E80 E82 E101 E113 E127 E130 E132
Cy.ENS ENS_PG NB Neu_A Neu_B Glia_1 Glia_2 Glia_3
C
Time
D
Marker Expression
TOP2A MKI67 TYMS CDH2 SOX10 SOX2 NPM1 ASCL1 DLL3 HAND2 TUBB3 UCHL1 PHOX2B ELAVL4 STMN2 ETV1 PENK NOS1 VIP NPY SCGN GAL BNC2 TAC1 CASZ1 SLC5A7 SST S100B ERBB3 PTPRZ1 BCAN APOE ENTPD2 RXRG TNC GFRA2 MPZ MAL DHH NTRK2 FGL2
Expression Level
Expression %
1%
25%
50%
75%
100%
E
S100B TUBB3 SOX10
PGP9.5 DAPI
Early
Arrowhead: S100B+TUBB3+SOX10+PGP9.5+; ENS_PG
F
S100B TUBB3 PGP9.5
nNOS DAPI
Late
Arrowhead: S100B-TUBB3+PGP9.5+nNOS+; Neu
Arrow: S100B+TUBB3-PGP9.5-nNOS-; Glia
G
Dim-1
Dim-2
H
ENDO
Cy.Endo
Art_1
Art_2
Ven_PG
Ven_1
Ven_2
Ven_3
Lym_1
Lym_2
I
Cell Proportion
J
Time
K
Marker Expression
TOP2A MKI67 TYMS CLDN5 PECAM1 CAV1 ESAM FLT1 PRND GJA4 CXCR4 PGF LXN CHST1 HEY1 GJA5 CXCL12 GRIA2 JAM2 APLNR PLVAP NOSTRIN RGCC ACKR1 ADGRG6 SELE SELP CD83 CA4 INMT LYVE1 PROX1 CCL21 TFF3 STAB2 SCG3 FOXC2 GATA2 DPP4 CLU
L
Art Ven Lym Peri Im
PECAM1
VWF
POSTN
LYVE1
ACTA2
CSPG4
PTPRC
Expression Level
Expression %
M
VWF CD31 LYVE1 NG2 POSTN αSMA DAPI
Late
Arrow: VWF-CD31+LYVE1+NG2-POSTNloαSMA-; Lym
Arrowhead: VWF+CD31+LYVE1-NG2-POSTN-αSMA-; Art
Arrow: VWF+CD31+LYVE1-NG2-POSTN+αSMA-; Ven
Arrowhead: VWF-CD31-LYVE1-NG+POSTN-αSMA+; Peri
N
CD45 DAPI
Late
Arrowhead: CD45+; Im

### Figure S3

A

B

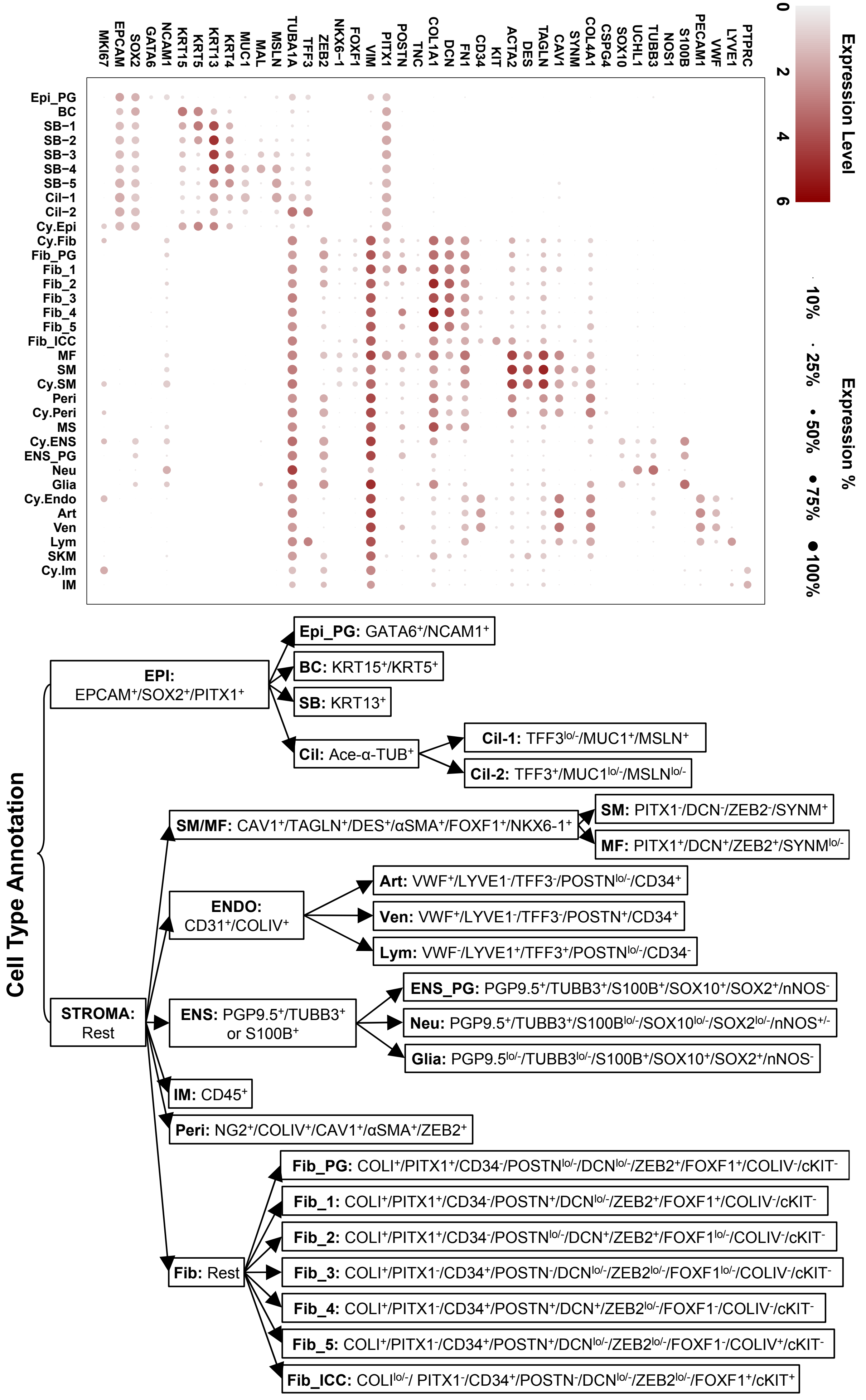
MKI67
EPCAM
SOX2
GATA6
NCAM1
KRT15
KRT5
KRT13
KRT4
MUC1
MAL
MSLN
TUBA1A
TFF3
ZEB2
NKX6-1
FOXF1
VIM
PITX1
POSTN
TNC
COL1A1
DCN
FN1
CD34
KIT
ACTA2
DES
TAGLN
CAV1
SYNM
COL4A1
CSPG4
SOX10
UCHL1
TUBB3
NOS1
S100B
PECAM1
VWF
LYVE1
PTPRC
Epi_PG
BC
SB-1
SB-2
SB-3
SB-4
SB-5
Cil-1
Cil-2
Cy.Epi
Cy.Fib
Fib_PG
Fib_1
Fib_2
Fib_3
Fib_4
Fib_5
Fib_ICC
MF
SM
Cy.SM
Peri
Cy.Peri
MS
Cy.ENS
ENS_PG
Neu
Glia
Cy.Endo
Art
Ven
Lym
SKM
Cy.Im
IM
Expression Level
0
2
4
6
Expression %
10%
25%
50%
75%
100%
Cell Type Annotation
EPI: EPCAM+/SOX2+/PITX1+
Epi_PG: GATA6+/NCAM1+
BC: KRT15+/KRT5+
SB: KRT13+
Cil: Ace-α-TUB+
Cil-1: TFF3lo/-/MUC1+/MSLN+
Cil-2: TFF3+/MUC1lo/-/MSLNlo/-
STROMA: Rest
SM/MF: CAV1+/TAGLN+/DES+/αSMA+/FOXF1+/NKX6-1+
SM: PITX1-/DCN-/ZEB2-/SYNM+
MF: PITX1+/DCN+/ZEB2+/SYNMlo/-
ENDO: CD31+/COLIV+
Art: VWF+/LYVE1-/TFF3-/POSTNlo/-/CD34+
Ven: VWF+/LYVE1-/TFF3-/POSTN+/CD34+
Lym: VWF-/LYVE1+/TFF3+/POSTNlo/-/CD34-
ENS: PGP9.5+/TUBB3+ or S100B+
ENS_PG: PGP9.5+/TUBB3+/S100B+/SOX10+/SOX2+/nNOS-
Neu: PGP9.5+/TUBB3+/S100Blo/-/SOX10lo/-/SOX2lo/-/nNOS+/-
Glia: PGP9.5lo/-/TUBB3lo/-/S100B+/SOX10+/SOX2+/nNOS-
IM: CD45+
Peri: NG2+/COLIV+/CAV1+/αSMA+/ZEB2+
Fib: Rest
Fib_PG: COLI+/PITX1+/CD34-/POSTNlo/-/DCNlo/-/ZEB2+/FOXF1+/COLIV-/cKIT-
Fib_1: COLI+/PITX1+/CD34-/POSTN+/DCNlo/-/ZEB2+/FOXF1+/COLIV-/cKIT-
Fib_2: COLI+/PITX1+/CD34-/POSTNlo/-/DCN+/ZEB2+/FOXF1lo/-/COLIV-/cKIT-
Fib_3: COLI+/PITX1-/CD34+/POSTN-/DCNlo/-/ZEB2lo/-/FOXF1lo/-/COLIV-/cKIT-
Fib_4: COLI+/PITX1-/CD34+/POSTN+/DCN+/ZEB2lo/-/FOXF1-/COLIV-/cKIT-
Fib_5: COLI+/PITX1-/CD34+/POSTN+/DCNlo/-/ZEB2lo/-/FOXF1-/COLIV+/cKIT-
Fib_ICC: COLIlo/-/ PITX1-/CD34+/POSTN-/DCNlo/-/ZEB2lo/-/FOXF1+/cKIT+

### Figure S4

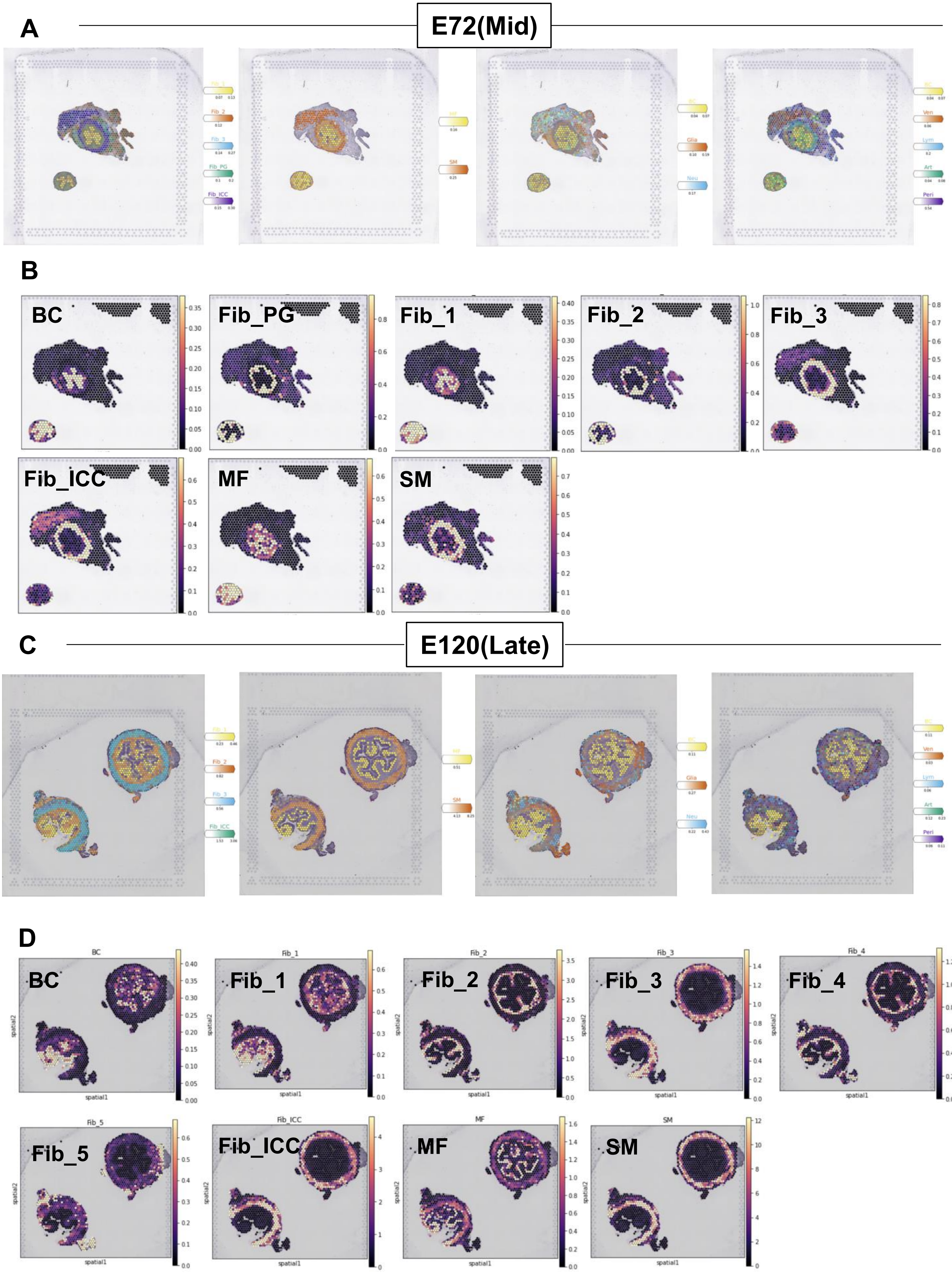
A
E72(Mid)
B
BC
Fib_PG
Fib_1
Fib_2
Fib_3
Fib_ICC
MF
SM
C
E120(Late)
D
BC
Fib_1
Fib_2
Fib_3
Fib_4
Fib_5
Fib_ICC
MF
SM

Figure S5

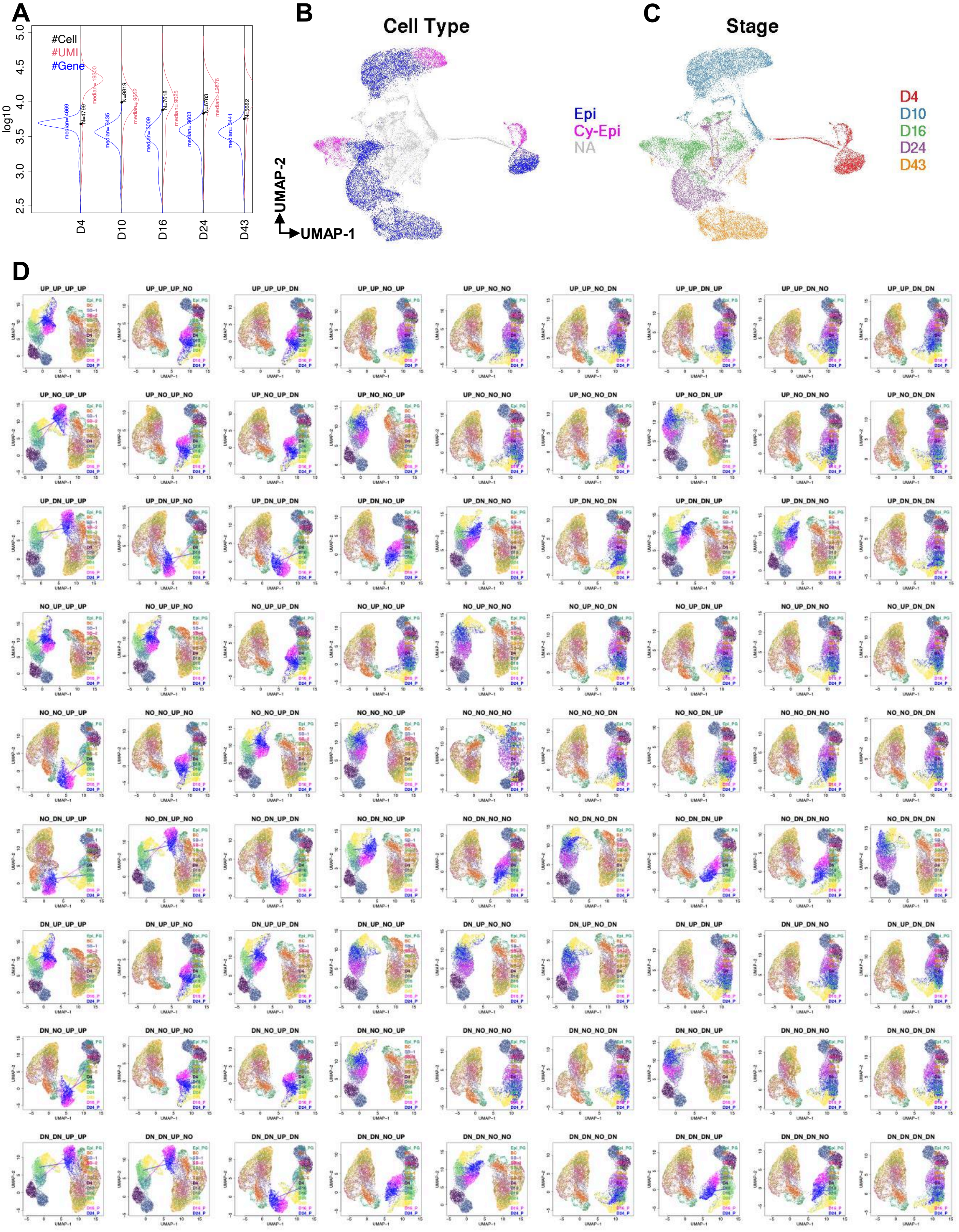
A
#Cell
#UMI
#Gene
log10
D4
D10
D16
D24
D43
B
Cell Type
Epi
Cy-Epi
NA
UMAP-2
UMAP-1
C
Stage
D4
D10
D16
D24
D43
D
UP_UP_UP_UP
UP_UP_UP_NO
UP_UP_UP_DN
UP_UP_NO_UP
UP_UP_NO_NO
UP_UP_NO_DN
UP_UP_DN_UP
UP_UP_DN_NO
UP_UP_DN_DN
UP_NO_UP_UP
UP_NO_UP_NO
UP_NO_UP_DN
UP_NO_NO_UP
UP_NO_NO_NO
UP_NO_NO_DN
UP_NO_DN_UP
UP_NO_DN_NO
UP_NO_DN_DN
UP_DN_UP_UP
UP_DN_UP_NO
UP_DN_UP_DN
UP_DN_NO_UP
UP_DN_NO_NO
UP_DN_NO_DN
UP_DN_DN_UP
UP_DN_DN_NO
UP_DN_DN_DN
NO_UP_UP_UP
NO_UP_UP_NO
NO_UP_UP_DN
NO_UP_NO_UP
NO_UP_NO_NO
NO_UP_NO_DN
NO_UP_DN_UP
NO_UP_DN_NO
NO_UP_DN_DN
NO_NO_UP_UP
NO_NO_UP_NO
NO_NO_UP_DN
NO_NO_NO_UP
NO_NO_NO_NO
NO_NO_NO_DN
NO_NO_DN_UP
NO_NO_DN_NO
NO_NO_DN_DN
NO_DN_UP_UP
NO_DN_UP_NO
NO_DN_UP_DN
NO_DN_NO_UP
NO_DN_NO_NO
NO_DN_NO_DN
NO_DN_DN_UP
NO_DN_DN_NO
NO_DN_DN_DN
DN_UP_UP_UP
DN_UP_UP_NO
DN_UP_UP_DN
DN_UP_NO_UP
DN_UP_NO_NO
DN_UP_NO_DN
DN_UP_DN_UP
DN_UP_DN_NO
DN_UP_DN_DN
DN_NO_UP_UP
DN_NO_UP_NO
DN_NO_UP_DN
DN_NO_NO_UP
DN_NO_NO_NO
DN_NO_NO_DN
DN_NO_DN_UP
DN_NO_DN_NO
DN_NO_DN_DN
DN_DN_UP_UP
DN_DN_UP_NO
DN_DN_UP_DN
DN_DN_NO_UP
DN_DN_NO_NO
DN_DN_NO_DN
DN_DN_DN_UP
DN_DN_DN_NO
DN_DN_DN_DN

### Figure S6

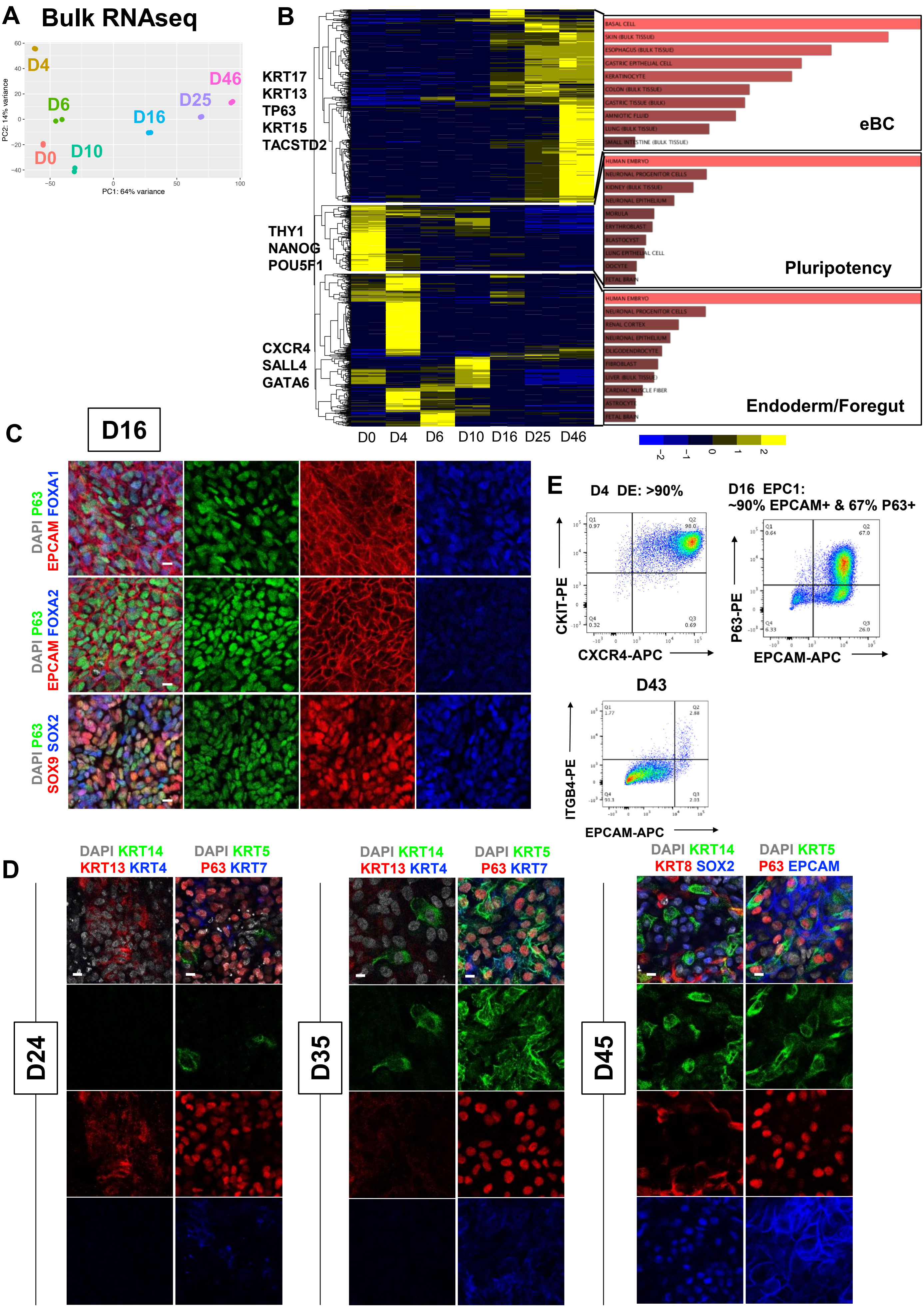
A Bulk RNAseq
D4
D6
D0
D10
D16
D25
D46
PC1: 64% variance
PC2: 14% variance
B
KRT17
KRT13
TP63
KRT15
TACSTD2
THY1
NANOG
POU5F1
CXCR4
SALL4
GATA6
D0 D4 D6 D10 D16 D25 D46
BASAL CELL
SKIN (BULK TISSUE)
ESOPHAGUS (BULK TISSUE)
GASTRIC EPITHELIAL CELL
KERATINOCYTE
COLON (BULK TISSUE)
GASTRIC TISSUE (BULK)
AMNIOTIC FLUID
LUNG (BULK TISSUE)
SMALL INTESTINE (BULK TISSUE)
eBC
HUMAN EMBRYO
NEURONAL PROGENITOR CELLS
KIDNEY (BULK TISSUE)
NEURONAL EPITHELIUM
MORULA
ERYTHROBLAST
BLASTOCYST
LUNG EPITHELIAL CELL
OOCYTE
FETAL BRAIN
Pluripotency
HUMAN EMBRYO
NEURONAL PROGENITOR CELLS
RENAL CORTEX
NEURONAL EPITHELIUM
OLIGODENDROCYTE
FIBROBLAST
LIVER (BULK TISSUE)
CARDIAC MUSCLE FIBER
ASTROCYTE
FETAL BRAIN
Endoderm/Foregut
C
D16
DAPI P63 EPCAM FOXA1
DAPI P63 EPCAM FOXA2
DAPI P63 SOX9 SOX2
E
D4 DE: >90%
CKIT-PE
CXCR4-APC
D16 EPC1:
~90% EPCAM+ & 67% P63+
P63-PE
EPCAM-APC
D43
ITGB4-PE
EPCAM-APC
D
DAPI KRT14 KRT13 KRT4
DAPI KRT5 P63 KRT7
DAPI KRT14 KRT8 SOX2
DAPI KRT5 P63 EPCAM
D24
D35
D45

### Figure S7
